# Conservation of molecular responses upon viral infection in the non-vascular plant *Marchantia polymorpha*

**DOI:** 10.1101/2023.11.06.565823

**Authors:** Eric Ros-Moner, Tamara Jiménez-Góngora, Luis Villar-Martín, Lana Vogrinec, Víctor M. González-Miguel, Denis Kutnjak, Ignacio Rubio-Somoza

**Affiliations:** Molecular Reprogramming and Evolution (MoRE) Laboratory, Centre for Research in Agricultural Genomics (CRAG), CSIC-IRTA-UAB-UB, Campus UAB-Edifci CRAG. 08193 Cerdanyola del Vallés, Spain; National Institute of Biology, Department of Biotechnology and Systems Biology, Ljubljana, Slovenia; Jožef Stefan International Postgraduate School, Ljubljana, Slovenia; Data Analysis area, Bioinformatics Core Unit, Centre for Research in Agricultural Genomics (CRAG), CSIC-IRTA-UAB-UB, Campus UAB-Edifci CRAG. 08193 Cerdanyola del Vallés, Spain; Consejo Superior de Investigaciones Científicas (CSIC), Barcelona 08001, Spain

## Abstract

After plants transitioned from water to land around 450 million years ago, they faced novel pathogenic microbes. Their colonization of diverse habitats was driven by anatomical innovations like roots, stomata, and vascular tissue, which became central to plant-microbe interactions. However, the impact of these innovations on plant immunity and pathogen infection strategies remains poorly understood. Here, we explore plant-virus interactions in the bryophyte *Marchantia polymorpha* to gain insights into the evolution of these relationships. Virome analysis reveals that *Marchantia* is predominantly associated with RNA viruses. Comparative studies with tobacco mosaic virus (TMV) show that *Marchantia* shares core defense responses with vascular plants but also exhibits unique features, such as a sustained wound response preventing viral spread. Additionally, general defense responses in Marchantia are equivalent to those restricted to vascular tissues in Nicotiana, suggesting that evolutionary acquisition of developmental innovations results in re-routing of defense responses in vascular plants.

## Introduction

It is estimated that plants developed a functional vascular system through repurposing of pre-existing genes around 420 million years ago, which facilitated the growth of larger plants by providing mechanical support and enabling long distant transport of water, nutrients, and signalling molecules (Bowles et al., 2022, Lucas et al., 2013). Viruses are obligate parasites that can infect virtually any living organism hijacking central host cellular functions to replicate and spread within and intra hosts. Coincidentally with land colonization, the repertoire of viruses (virome) associated to plants enlarged and diversified both in nature and infection strategies. Although our current knowledge lacks information on virome composition at intermediately divergent plant lineages, it appears that viruses with large DNA genomes majorly infecting unicellular chlorophyte algae were displaced by an increasing population of viruses with RNA and small DNA genomes found in vascular plants (Mushegian et al., 2016). Plant vasculature has become central for the success of viral infections in tracheophytes, not only serving as fast track for viral systemic spread, but also providing a unique cellular environment for some viruses which replication is restricted to that tissue (Kappagantu et al., 2020). Accordingly, vascular cells are thought to present specific responses to viral infection when compared to other leaf cell-types (Collum and Culver, 2017).

Our current understanding of the molecular events governing plant-virus interactions is vastly shaped by studies focusing on vascular plants, mainly crops. Plants respond to viral infections via at least three different but interconnected defense mechanisms (Mandadi and Scholthof, 2013). Firstly, plants have an extracellular surveillance system that monitors the presence of pathogens through the recognition of highly conserved microbe patterns dubbed as Pattern-associated molecular patterns (PAMPs), such as viral derived double-strand RNAs (dsRNAS), the bacterial flagellin or the fungal chitin. Detection of those molecules by specialized surface-localized receptor kinases (Pattern recognition receptors, PRRs), triggers a first layer of defense called PAMP-triggered immunity (PTI). PRRs expression is often restricted to focal points for microbial interaction and invasion (Beck et al., 2014, Yotsui et al., 2023). Two additional surveillance mechanisms monitor the intracellular presence of microbe-derived molecules. Highly stable dsRNAs produced by viruses during their life cycle, independently of the DNA or RNA nature of their genomes, are detected by the RNA silencing machinery which constitutes a conserved antiviral defense mechanism both in plants and animals (Ding et al., 2018, Ding and Voinnet, 2007). Additionally, RNA silencing machinery also recognizes dsRNA generated from the conversion of exogenous and endogenous single strand RNAs (ssRNA) by RNA-dependent RNA polymerases (RDRs). RDR1 and RDR6 have been involved in antiviral defense in *Arabidopsis thaliana* and *Nicotiana tabacum* (Garcia-Ruiz et al., 2010, Rakhshandehroo et al., 2017, Ying et al., 2010), even though *Nicotiana benthamiana* has only a functional RDR6 (Bally et al., 2015). Upon recognition, viral dsRNAs are trimmed in small RNAs (visRNAs) of 21 to 24 nucleotides in length by members of the DICER-like family, mainly DCL2, DCL3 and DCL4 in Arabidopsis and Nicotiana (Garcia-Ruiz et al., 2010, Lewsey and Carr, 2009). The resulting visRNA duplexes are subsequently loaded into proteins from the ARGONAUTE family, which are the core elements of the RNA-induced silencing complex (RiSC), and used to scan the intracellular space for the source of dsRNA precursors in a sequence complementary manner abrogating viral infection (Lopez-Gomollon and Baulcombe, 2022). In turn, viruses produce countermeasures, known as silencing suppressors, targeted to interfere with different steps of the host RNA silencing machinery (Lopez-Gomollon and Baulcombe, 2022). In response to these silencing suppressors, host micro RNAs (miRNA, one of the two main classes of endogenous sRNAs in plants) are no longer functional, allowing the translation of their targets and licensing a new layer of defense. Among the miRNA targets involved in such counter-counter measure, members of the Nucleotide-binding domain leucine-rich repeat (NLR) family are central. Thus, it has been suggested that host miRNAs might work as sensors for the presence of pathogen-derived silencing suppressors enabling a rapid host defense response based on the activation of otherwise silenced miRNA targets (Vasseur et al., 2022, Silvestri et al., 2024). NLRs are core elements of the third intracellular surveillance system for directly monitoring the intracellular presence of pathogen-derived molecules, such as the prototypical case of the tobacco mosaic virus (TMV) replicase by the gene *N* in tobacco (Erickson et al., 1999), or their impact on host cellular homeostasis. NLR-activation triggers the so-called Effector-triggered immunity (ETI), which normally leads to a hypersensitive response (HR) culminating in cell death. Those three defense systems, PTI, RNA silencing and ETI largely rely on the action of the plant hormone Salicylic acid (SA). PTI and ETI responses encompass activation of SA signalling (Pruitt et al., 2021) while SA activates RDR1 in Arabidopsis and Nicotiana (Lee et al., 2016, Ying et al., 2010, Yu et al., 2003). Additionally, RDR1 expression is also induced by treatment with jasmonates (Liu et al., 2009, Xu et al., 2013), another central hormone in plant defense. Many of those immunity components and networks were already present in the last common land plant ancestor, while others have been acquired alongside the increasing plant cellular and anatomical complexity (de Vries et al., 2018, Belanger et al., 2023, Chia et al., 2024b).

A central question in plant-microbe interactions is how the evolution of different plant developmental trajectories, including the acquisition of major innovations, impacts host defense and pathogen infection strategies. In animals, comparative studies have proved very informative about the origin and diversification of their innate immune responses against virus (Iwama and Moran, 2023). Likewise, comparative studies including non-vascular plants, like the liverwort *Marchantia polymorpha*, have been instrumental in understanding the evolution of the interactions between plants and pathogens other than viruses (Carella et al., 2018, Gimenez-Ibanez et al., 2019, Redkar et al., 2022). Comparative studies in plants have traditionally relied on both model plant species and microbes, despite the lack of prior experimental evidence for those interactions to occur in nature. Incorporating bryophytes into those comparative studies not only provides important information on how plant immune programs may have evolved, but it also bears the potential to determine the contribution of developmental innovations, such as roots, stomata or vasculature, to their diversification.

To enable comparative studies towards understanding the evolution of plant-virus interactions, we here characterized the molecular interplay between the non-vascular liverwort *Marchantia polymorpha* and viruses. We firstly defined the composition of the virome of Marchantia plants living in the wild and used that information to select a virus with similar genomic features to those found in nature for further molecular characterization, and which could be tagged with a fluorescent protein that enables monitoring of infection. Additionally, the selected virus should be able to infect a broad range of hosts, in which their molecular interactions have been studied. Following those criteria, we focused on the first ever isolated virus, the tobacco mosaic virus (TMV). TMV is a ssRNA virus from the Tobamovirus genus showing a broad host range in nature (Zamfir et al., 2023). Additionally, TMV has been extensively used for studying plant-virus interactions in a wealth of plant species, including several Solanaceae species, like tomato and tobacco, and the model plant *Arabidopsis thaliana*, which provide strong foundation for comparative studies. TMV infects through wound entry points from where it moves cell-to-cell via plasmodesmata, until it reaches the vascular system for systemic infection of distal parts of the plant following the stream of photoassimilates (Liu and Nelson, 2013).

Our results show that *Marchantia polymorpha* is associated with RNA viruses in natural populations that can replicate in lab grown Marchantia plants. Additionally, TMV can replicate in Marchantia plants triggering a conserved transcriptional reprogramming encompassing increased wounding and PTI responses in detriment of photosynthesis and cell cycle progression and leading to changes in life-history traits, such as aging and senescence. Induction of a wealth of NLR proteins and core elements from the RNA silencing machinery were also observed, including DCL4 that, in absence of a DCL2 gene in the Marchantia genome (Belanger et al., 2023), seems to be the main antiviral DCL protein according to the predominant 21 nuleotides in length presented by TMV-derived vsRNAs. Noteworthy, the general response of the RNA silencing machinery observed in Marchantia upon TMV infection is equivalent to that found specifically in vascular tissues of *Nicotiana benthamiana*, suggesting an evolutionary anatomical rerouting of antiviral defense towards newly acquired focal points for viral infection.

We additionally identified a conserved host interactor with the TMV silencing suppressor p126 that triggers cross-species defense against TMV.

## Results

### *Marchantia polymorpha* is associated to RNA viruses in nature

To identify viruses associated with *Marchantia polymorpha* in nature, we performed a virome analysis of two composite plant samples including 21 individuals growing naturally, not cultivated, in 5 different locations spanning 4 different countries and accessible to possible viral vectors. First included Marchantia individuals obtained from three open air botanical gardens throughout Europe: Tübingen (Germany), Bern (Switzerland) and Cambridge (UK). The second encompassed individuals growing open air at the botanical garden in Osnabrück (Germany) and individuals growing in urban sites in Ljubljana (Slovenia). In addition to the expected background of bacteriophage sequences, we detected several known viruses, including some capable of infecting or being associated with plants, fungi or invertebrates (Supplementary Data 1). Putatively new virus-like sequences, possibly associated with *M. polymorpha*, were selected for further analyses from the first dataset, based on i) their relatively high average sequencing depths and ii) similarity to viruses previously associated with eukaryotic organisms, including plants and invertebrates (Supplementary Data 2). Two previously described and three newly identified viruses were hence selected for further RT-qPCR confirmation of their association with the different *M. polymorpha* individuals, along with plants from the Tak-1 reference accession grown in sterile conditions. Specific primers to detect Ligustrum necrotic ringspot virus (LNRV, known to infect plants), Hubei picorna-like virus 51, (detected in a metagenomics study and putatively associated with diverse sample types by datamining – see results below) a newly identified picorna-like virus (which we named Tübingen picorna-like virus 1, with average mapping depth of 2230x and contig length of 7801 nts), a new tombus-like virus (Tübingen tombus-like virus 1, with average mapping depth of 218x and contig length of 4521 nts), and a new sobemo-like virus (Bern sobemo-like virus 1, with average mapping depth of 88x and contig length of 2569 nts) were used for these analysis. While none of those viruses were detected in negative control Tak-1 plants, individuals from Tübingen were associated with all tested viruses but the Bern sobemo-like virus 1, which in turn was present in individuals from Bern along with Tübingen picorna-like virus 1. Individuals from the Cambridge population were associated with Hubei picorna-like virus 51 and Tübingen picorna-like virus 1 (Supplementary Data 3). Further, we used a datamining approach to search for the presence of conserved RdRp domain fingerprints (palmprints) of the selected viruses in global short reads sequence datasets using Serratus infrastructure (Edgar et al., 2022). The results of the analysis suggested that palmprint RdRp sequences with high similarity to those of queried viruses, can be associated with diverse sample types, mostly including environmental samples, such as wastewater, water, rhizosphere, soil and plants (Supplementary Data 4).

Given that Tübingen picorna-like virus 1 was present in all tested populations from the first sampling, we decided to further characterize its genomic features and interaction with *Marchantia polymorpha*. Indeed, Tübingen picorna-like virus 1 had the highest mapping depth of all contigs in our virome analysis and showed a similar genome structure as other unclassified picorna-like viruses (Shi et al., 2016), consisting of two predicted genome open reading frames (ORFs), first containing domains with homology to superfamily 3 helicase, peptidase C3 and RNA-dependent RNA polymerase (RdRp) and second three domains with homology to a coat protein with 8-stranded jelly roll β-barrel motif (Figure 1A). Phylogenetic analysis (Figure 1B) of the RdRp protein sequence and its closest NCBI GenBank homologues showed high similarity (80% identity of aligned sequences) of this virus isolate with a recently reported plant associated picorna-like virus (Trichosanthes kirilowii picorna-like virus, QKK82970.1). Much lower identities were observed for Tübingen picorna-like virus 1 (29-42%) and other viruses associated with diverse sources (plants, gastropods, water, crayfish, artropods and bird feces). However, given that they were discovered in metagenomics studies their association with these hosts is only speculative. Thus, we performed and additional phylogenetic analysis, comparing the RdRp sequence of Tübingen picorna-like virus 1 with most similar RdRp sequences from RNA Viruses in Metatranscriptomes (RVMT) database (https://riboviria.org/) (Neri et al., 2022). The analysis revealed high similarity of the viral RdRp to several sequences included in family-level cluster f.0032 with order *Picornavirales* (Supplementary Fig. 1A). Even if this cluster contains diverse representatives, none of those has confirmed host associations to the date (Supplementary Data 5). Similarly, as found in Serratus analysis, sequences most similar to investigated virus were associated, e.g., with wastewater, wetlands, and peat soil samples (Supplementary Fig. 1b, Supplementary Data 5).

**Figure 1.**
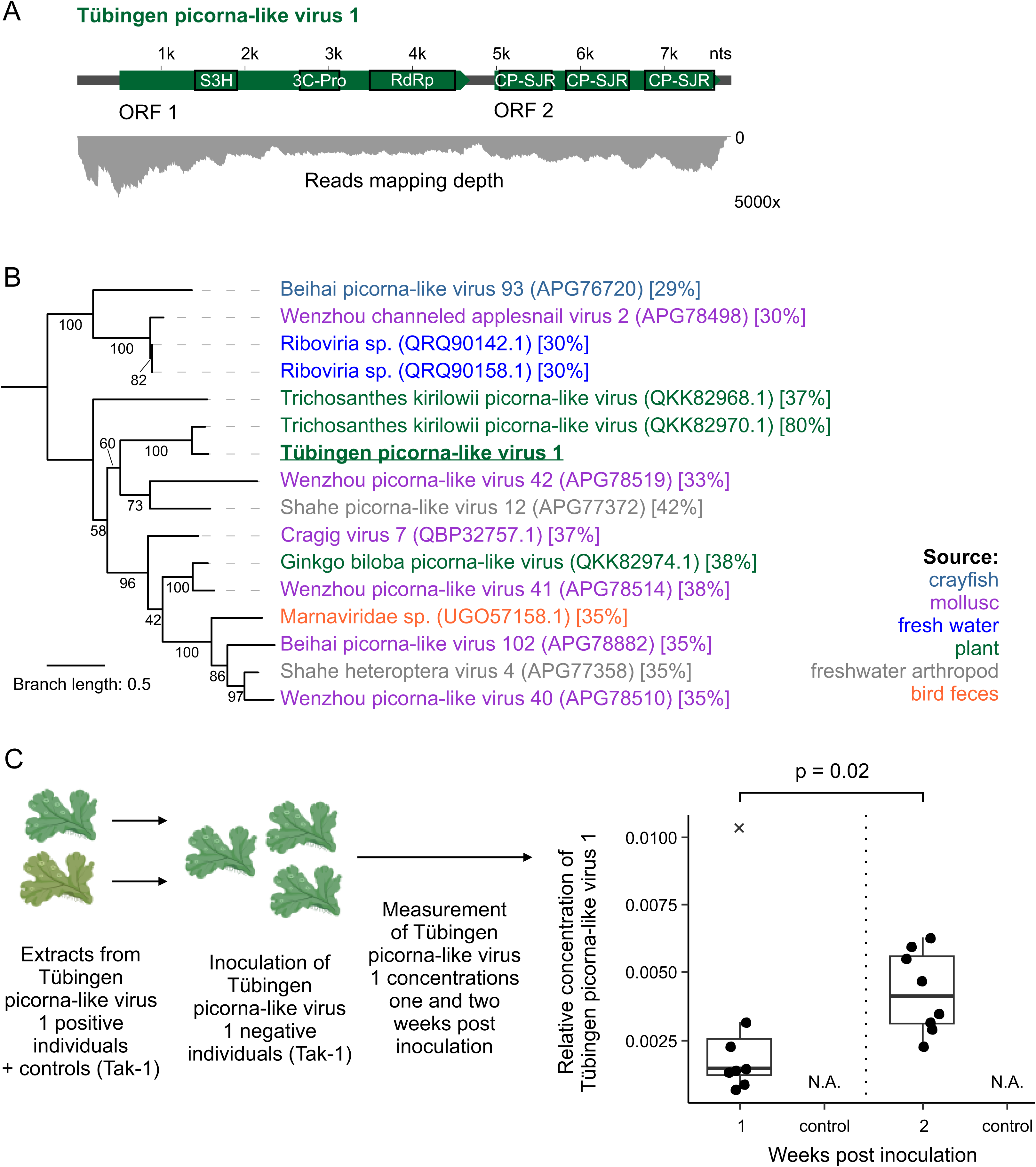
Marchantia polymorpha is infected by RNA viruses. (A) Genome structure of Tübingen picorna-like virus 1, newly discovered in composite *M. polymorpha* sample with two predicted open reading frames (ORFs) and corresponding average reads mapping depth in sequenced composite sample. Black empty boxes represent predicted protein domains detected by InterProScan: S3H – superfamily 3 helicase, 3C-Pro – peptidase C3, RdRp – RNA-dependent RNA polymerase and CP-SJR – coat protein with 8-stranded jelly roll β-barrel motif. (B) Maximum likelihood phylogenetic tree obtained based on the alignment of the conserved part of RdRd of Tübingen picorna-like virus 1 and fifteen other most similar viruses. The tree was midpoint rooted. Coloured virus names correspond to the sources associated with a particular virus, according to the legend on the right side of the panel. Numbers next to the branches represent statistical support (%) of the clades according to the ultrafast bootstrap analysis. Branch length represents the average number of amino acid substitutions per site. Codes next to the taxon names in round brackets are the corresponding NCBI GenBank protein accession numbers. Next to them, fractions in square brackets represent % of identity of Tübingen picorna-like virus 1 to other individual taxons for the aligned part of the RdRp domain. (C) Results of the viral concentration measurements for *M. polymorpha* plants inoculated with Tübingen picorna-like virus 1 positive plant extracts. On the right, relative concentrations of Tübingen picorna-like virus 1 in inoculated plants one-and two-weeks post inoculation are shown; measurements of concentrations in individual plants (from 3 technical replicates each) are shown as dots (n=8 for both time points) and the box-whisker plots represent the distribution of values (showing median as a thick line, first and third quartile as boxes and minimum and maximum as whiskers) for the two time points (at 1 WPI, x represent an outlier measurement). On the top of the plot, the horizontal connecting line designates statistically significant difference in the means of the two distributions, with the corresponding p-value, according to the nonparametric Wilcoxon rank sum test (two-sided). Source data are provided as a Source data File. BIORENDER was used to prepare this figure.

Next, we tested whether the Tübingen picorna-like virus 1 was able to replicate in *M. polymorpha* by incubating two-weeks old Tak-1 individuals with extracts isolated from the Tübingen population, which contained the highest levels of Tübingen picorna-like virus 1, as determined by RT-qPCR analysis, or from Tak-1 plants grown in sterile conditions. Individual Marchantia plants were collected one and two weeks after incubation and the levels of viral accumulation were measured by RT-qPCR. Individuals exposed to extracts from virus-positive plants showed a trend of increased viral levels over time, while no amplification was observed in plants incubated with extracts from Tak-1 plants (Figure 1C). This indicated that Tübingen picorna-like virus 1 can replicate in Marchantia.

Collectively, those results show that *Marchantia polymorpha* can be associated with RNA viruses in nature.

Additionally, we performed data mining to search for possible DNA viruses associated with *Marchantia polymorpha*. Sequencing data from 74 biological samples (obtained from natural environments; (Beaulieu et al., 2023)) was analyzed for the presence of viral contigs. Analysis revealed that most of the examined samples contained virus-like sequences, with the majority exhibiting similarity to bacteriophages (Supplementary Data 6). Within the dataset we also found virus-like sequences with similarity to members of *Phycodnaviridae,* which can infect unicellular green algae. Since only few relatively short contig sequences for this group of viruses were identified in different datasets, and the data was primarily not produced with microbiome study in mind (e.g., no surface washing) the association with *M. polymorpha* with viruses from this taxon is dubious and shall be further explored in future studies.

### Tobacco mosaic virus replicates in *Marchantia polymorpha*

Since its identification in the 19^th^ century, tobacco mosaic virus has been extensively used for studying plant-virus interactions in several vascular plant species like Arabidopsis and Nicotiana (Scholthof, 2004). As a first step for comparative studies on plant-virus interactions between vascular and non-vascular plants, we aimed to infect *Marchantia polymorpha* plants with a GFP labelled clone of TMV (Peart et al., 2002). Sap from leaves of healthy and infected *Nicotiana benthamiana* plants were isolated and used to incubate two-week-old Tak-1 plants grown in sterile conditions. After incubation, Marchantia plants were transferred to soil and part of the remaining sap was mechanically inoculated with an abrasive (carborundrum) in basal leaves from 1 month old Nicotiana plants, as control for its infectivity. After two weeks, Nicotiana plants showed systemic GFP signal under UV light along with symptoms of infection (Supplementary Fig. 2), whereas Tak-1 plants displayed GFP signal in discrete infection foci that increased their size over time (Figure 2A, Supplementary Fig. 3), suggestive of viral replication. This was further corroborated by RT-qPCR amplification of pooled samples including four GFP positive individuals at the first (T1) and second weeks after incubation (T2), in comparison with mock and freshly incubated (T0) plants (Figure 2B, Supplementary Fig. 4). sRNA profiling in GFP positive areas from Marchantia thalli identified TMV-derived vsRNAs enriched in 21 nucleotides long fragments, consistent with DCL4-mediated processing, and mapping to the TMV-GFP sequence (Figure 2C, Supplementary Fig. 5).

**Figure 2.**
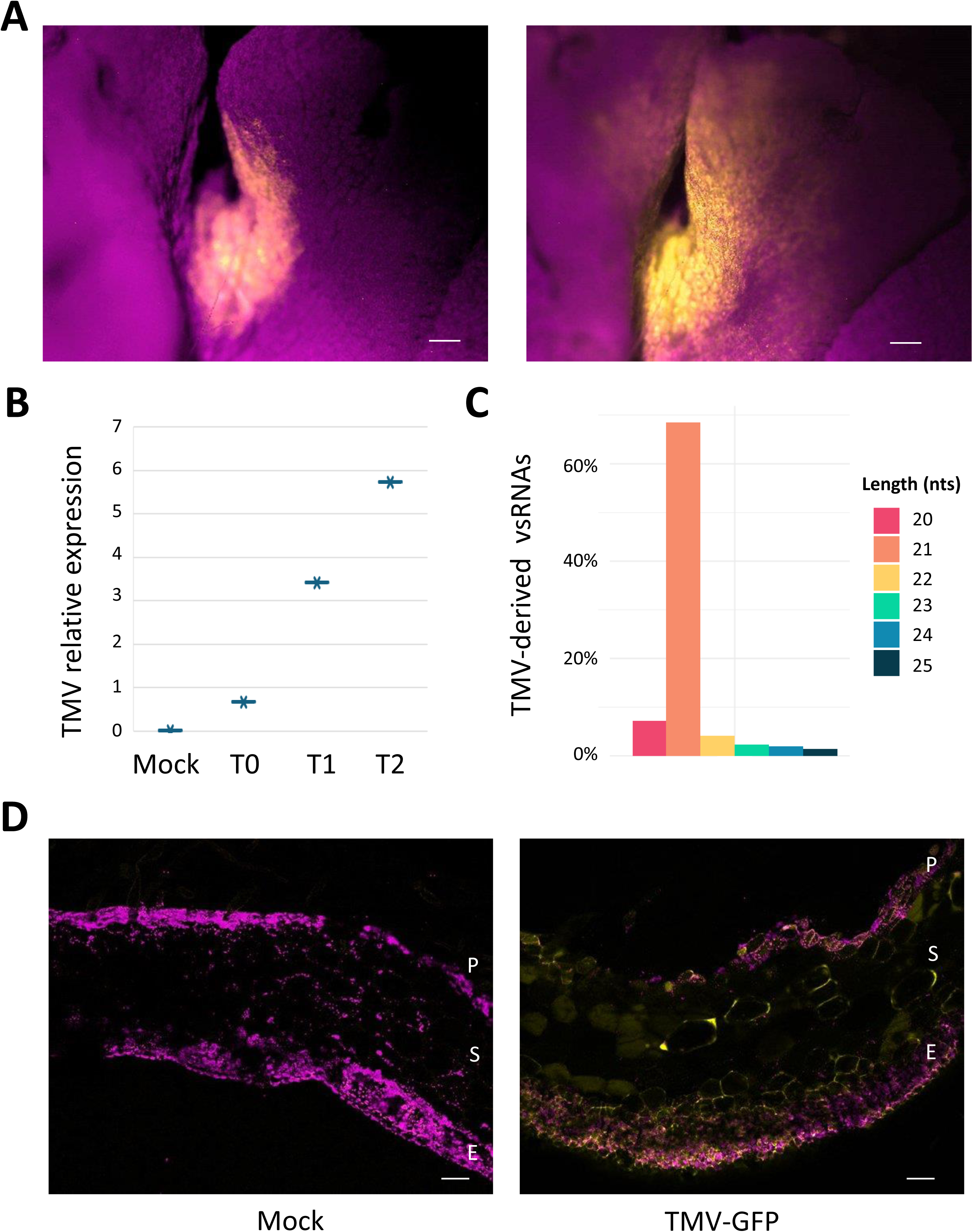
Tobacco mosaic virus replicates in Marchantia plants. (A) GFP fluorescent signal (in yellow) increases in thalli from 3-week-old plants after 1 week, (left panel) compared to thalli from 4 weeks old and 2-weeks infected plants (right panel). Both images show the same part or an infected plant under a magnification device under UV light. Magenta shows chlorophyll autofluorescence. White bar scale shows 100 µms. This experiment was repeated 4 times, and one additional replicate is shown in Supplementary Fig. 3. (B) TMV-GFP transcripts increase over time in infected plants. Pools of three infected plants were used for each time point. The result from 3 technical replicates is shown, with two complementary replicates showing the same trend in Supplementary Fig. 4. Mock, 4 weeks old plants incubated with sap from untreated plants. T0, 2 weeks old plants collected right after incubation with sap from infected plants. T1, 3 weeks old plants collected after 1 week from incubation. T2, 4 weeks old plants collected after 3 weeks from incubation. Expression of TMV is relative to the expression of Marchantia actine. Source data are provided as a Source data File. (C) Length distribution of TMV-derived vsRNAs populations found in thalli from 4 weeks old plants after 2 weeks of infection. Nts stans for nucleotides. (D) TMV-GFP infects all cell types. Transversal sections of an Marchantia thalli from a mock treated plant that was 4 weeks old (left) and equivalent area in a plant 2 weeks after infection (right) under confocal image. The pictures show the merge between green (in yellow, GFP-labelled virus) and red (in magenta, autofluorescence) channels. Merge was performed using Fiji. P stands for Photosynthetic region, S for storage region and E stands for lower epidermis. White bar scale shows 100 µm. This experiment was repeated 3 times showing similar results.

Histological sections of the infected area showed presence of the virus in all cell types, as inferred from GFP fluorescence and when compared to sections from mock treated plants, suggesting no preference for a specific cell-type during viral infection (Figure 2D).

Those results confirmed that TMV can replicate in *M. polymoprha* plants pointing to Marchantia-TMV as a suitable pathosystem for comparative studies.

### TMV infection triggers profound molecular reprogramming in *M. polymoprha*

To characterize the molecular reprogramming caused by TMV infection, we performed transcriptomic assays. TMV-GFP infected tissue from 4-week-old Marchantia plants (2 weeks after inoculation) was collected in four pools including samples from 5 plants each, along with the very same area from mock treated plants as control (Supplementary Fig. 6A). RNA was isolated and the presence of TMV was confirmed by RT-qPCR in each pool (Supplementary Fig. 6B). Principal component analysis (PCA, Supplementary Fig. 6C) of the transcriptome supported the RT-qPCR results showing a clear separation of infected and mock samples. Since separation among infected samples correlated with viral RT-qPCR levels, we selected the three replicates that had the highest levels of TMV and clustered together for further analysis. A total of 1879 genes were found to be up regulated while 1645 were reduced (padjust<0.05, Log2(fold-change) =>1 or <-1) in response to TMV infection (Supplementary Data 7). Consistently with an ongoing pathogenic interaction, we found several defense related marker genes among the differentially expressed genes (DEG). Members of the pathogenesis-related (PR) family are prime markers of immune responses in several plant species, including Marchantia (Carella et al., 2019, Gimenez-Ibanez et al., 2019) and we found that the expression of several homologues showed opposite expression profiles when compared to mock treated plants (Supplementary Table 1). Several PTI marker genes have been identified in Marchantia based on their orthology to those established in Arabidopsis (Gimenez-Ibanez et al., 2019) and their activation in the presence of bacterial and fungal extracts (Gimenez-Ibanez et al., 2019, Redkar et al., 2022). We observed that a set of those conserved PTI markers were also induced in response to TMV infection (Figure 3A). Additionally, a significant number of intracellular receptors from the NLR family displayed upregulated expression upon TMV infection (p < 6.245e-05 with those described by (Bowman et al., 2017); p < 1.308e-05 with those described by (Chia et al., 2024); Supplementary Table 2). Nevertheless, in line with the absence of TNL genes in Marchantia (Chia et al., 2024), none of those showed significant sequence similarity to the NLR N gene triggering ETI responses in TMV infected tobacco plants (Whitham et al., 1994). Likewise, several components from the RNA silencing machinery upregulated in tobacco plants upon TMV infection (Sheng et al., 2019), were also differentially expressed in response to TMV in Marchantia plants. The non-canonical MpDCL1b (Mp6g09830), MpDCL4 (Mp7g11720), and MpRDR6 (Mp8g14970) were upregulated, whereas MpDCL3 (Mp1g02840), MpAGO9 (Mp8g08610) and MpNRPE1b (Mp6g00230) were suppressed.

**Figure 3.**
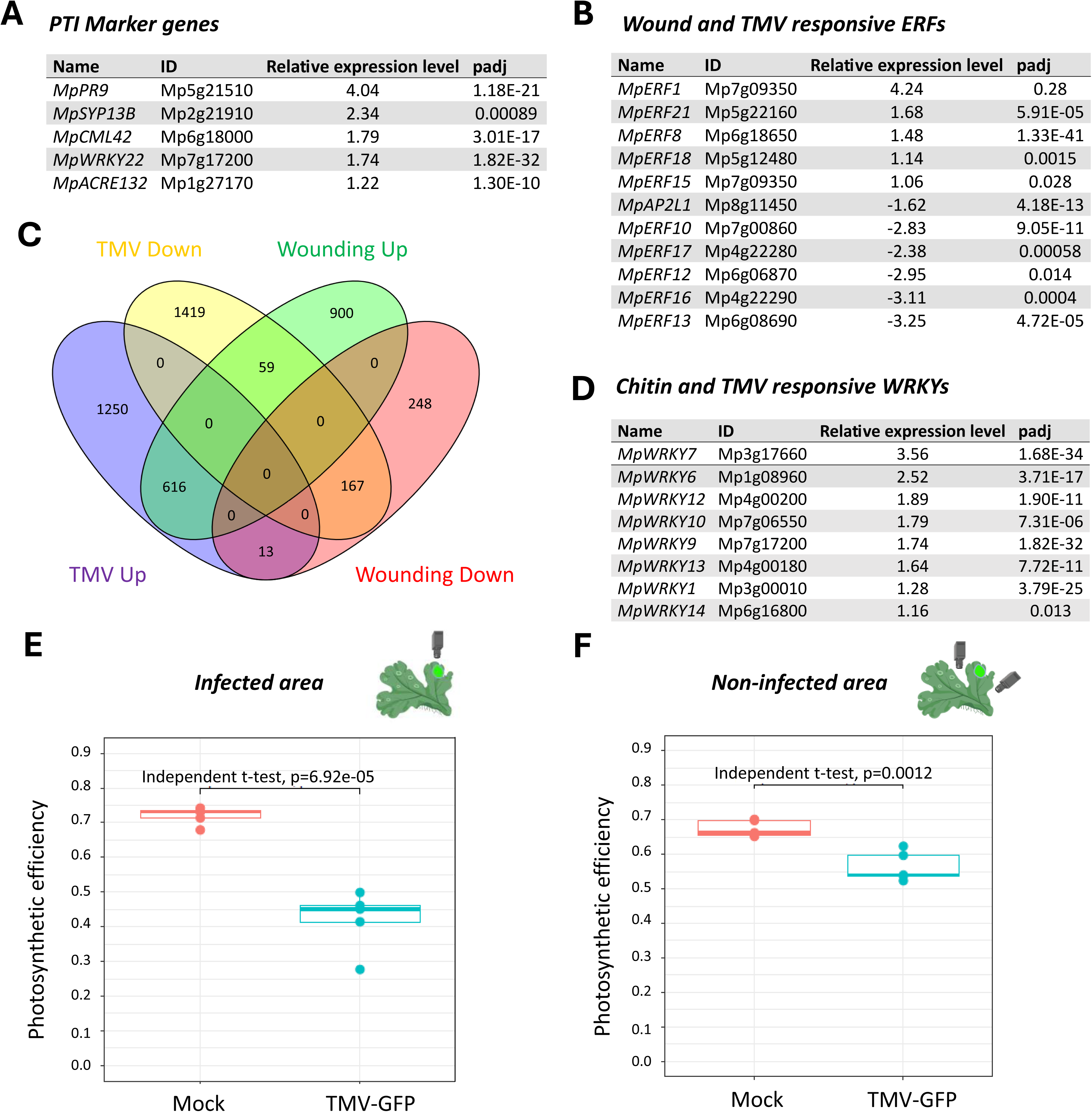
TMV triggers stress-related molecular reprogramming in Marchantia. (A) Set of common PTI marker genes responsive to bacterial and fungal extracts and TMV in infected areas in thalli from 4 weeks old plants after 2 weeks of infection compared to equivalent areas from mock treated plants. DESeq2 results using the default parameters (Wald test and Benjamini-Hochberg p adjustment method) (B) Set of wound and TMV responsive ERF transcription factors. DESeq2 results using the default parameters (Wald test and Benjamini-Hochberg p adjustment method) (C) Venn diagram showing that transcriptional reprogramming upon TMV infection largely overlaps with that found upon wounding. (D) Set of chitin and TMV responsive WRKY transcription factors in Marchantia. DESeq2 results using the default parameters (Wald test and Benjamini-Hochberg p adjustment method) (E) Photosynthetic efficiency (Fv/Fm) in infected areas in thalli from 4 weeks old plants after 2 weeks of infection compared to equivalent areas from mock treated plants. 5 biological and 10 technical replicates for each were assayed in both infected and mock treated plants. Statistical assessment was performed using a two-sided parametric T-test. Length of the whiskers: 1.5 x IQR. Central line: median. Box width: IQR (i.e. Q3-Q1). Source data are provided as a Source data File. Marchantia plants and cameras depicting measured areas come from Biorender. Light green area indicated TMV-GFP positive areas. (F) Photosynthetic efficiency (Fv/Fm) in non-infected areas, in thalli from 4 weeks old plants after 2 weeks of infection, compared to equivalent areas from mock treated plants. 5 biological and 10 technical replicates for each were assayed in both infected and mock treated plants. Statistical assessment was performed using a two-sided parametric T-test. Length of the whiskers: 1.5 x IQR. Central line: median. Box width: IQR (i.e. Q3-Q1). Source data are provided as a Source data File. Marchantia plants and cameras depicting measured areas come from Biorender. Light green area indicated TMV-GFP positive areas.

Gene Ontology (GO) analysis of the upregulated genes revealed an enrichment (padjust<0.05) of biological processes related to reactive oxygen species, wounding, light intensity, heat responses and responses to fungus and the presence of their constituent chitin (Supplementary Data 8, Supplementary Fig. 7A). Photosynthesis, DNA packaging and cell cycle, microtubule organization or regulation of salicylic acid metabolism were among the downregulated processes (padjust<0.05; Supplementary Data 9, Supplementary Fig. 7B).

Notably, TMV is mainly transmitted through wounds and wounding response was one of the processes to which upregulated genes belonged in the GO analysis, which prompted us analysing this response. ERF transcription factors are central to trigger wounding responses in vascular plants such as *A. thaliana* (Heyman et al., 2018) with a conserved set of ERF transcription factors being also recently described to respond to wounding in *Marchantia polymorpha* plants (Liang et al., 2022). A subset of these wounding responsive ERF transcription factors showed the same expression pattern upon TMV infection as that observed upon intensive wounding (Figure 3B, (Liang et al., 2022). To determine the whole transcriptional overlap between wounding and TMV responses, we reanalysed published tissue incision transcriptomic responses (Liang et al., 2022) using the same pipeline we used for the TMV samples. Comparison of both datasets showed a significant overlap both in up-regulated (593 genes; p<6.631e-235) and down-regulated genes (162 genes; p<2.137e-70; Figure 3C), suggesting that TMV infection might contribute to a sustained wound response in Marchantia.

Chitin is one of the most abundant polysaccharides in nature and it is a primary component of cell walls in fungi and exoskeletons of arthropods such as insects, which in turn can work as viral vectors in a range of plants species (Campbell, 1996). According to the GO enrichment in response to chitin and fungi, and besides the fungal responsive genes described above as part of a conserved PTI response, we also observed upregulation of a set of fungal responsive WRKY transcription factors (Figure 3D; (Redkar et al., 2022, Yotsui et al., 2023).

Photosynthetic deficiency is a hallmark for several stress responses, including those involving viral infections. Accordingly, downregulation of transcripts involved in photosynthesis was one of the main processes affected in our GO study as consequence of TMV infection. We determined the photosynthetic efficiency in TMV infected areas from 4-weeks old Tak-1 plants after 2-weeks of infection along with the very same area in mock treated plants. TMV infected areas showed a significant reduction in photosynthetic competency when compared to similar areas from mock treated plants (Figure 3E). Notably, parts of those very same plants distant to the infection foci, showed also reduced photosynthetic capability, although to a lesser extent (Figure 3F). Those results suggest that TMV responses might be systemic in *Marchantia polymorpha* plants.

Overall, those results show that TMV engages in a compatible interaction with *Marchantia polymorpha*.

### Salicylic and OPDA-dependent signaling pathways are active at TMV infected areas

The plant hormones Salicylic acid (SA) and Jasmonic acid (JA) are central in defense responses against most pathogens of vascular plants, acting both in a synergistic and antagonistic manner. Whereas the SA biosynthesis and signalling pathway are mostly conserved in *Marchantia polymoprha* (Jia et al., 2023), the active molecules binding to the COI1 receptor and triggering jasmonate signaling are a group of long chain polyunsaturated fatty acids, including dinor-OPDA, instead of JA-Ile (Kneeshaw et al., 2022, Monte et al., 2018). Therefore, we compared the SA and jasmonate-related transcriptomic responses with those found during TMV infection. To that end, we used the time course datasets obtained on incubation of Marchantia with 1mM SA during 2, 6 and 12 hours (Jeon et al., 2022), SA concentration shown to promote defense against bacterial and to increase susceptibility to fungal pathogens (Jeon et al., 2022, Matsui et al., 2020). Comparison of the DEGs identified on SA treatment showed a significant overlap with those responding to TMV infection (Figure 4A; Supplementary Fig. 8). A total of 225 genes were commonly up-regulated by 2 hours SA treatment or TMV (p<1.799e-100), 306 (p < 1.382e-172) were shared after 6 hours treatment, and 92 (p < 3.580e-47) by 12 hours of SA incubation. Likewise, TMV infection shared 47 down regulated genes (p<1.618e-14) with 2 hours, 38 (p < 2.897e-13) with the 6 hours, and 17 genes with 12 hours incubation samples (p < 1.720e-05), showing that TMV infection activates SA-dependent transcriptional responses.

**Figure 4.**
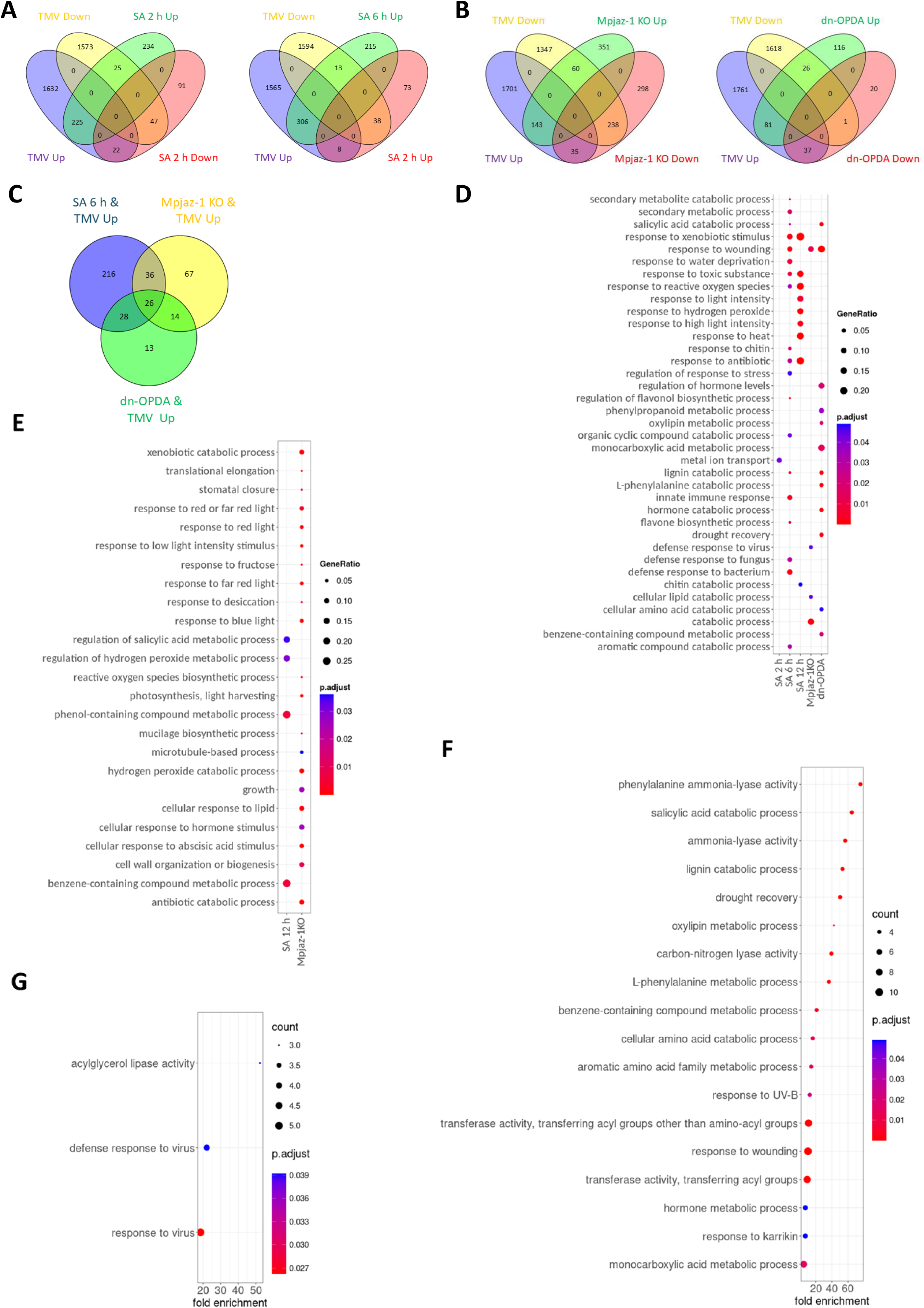
Salicylic and jasmonate signalling pathways are activated during Marchantia-TMV interaction. (A) Venn diagrams showing the overlap between the transcriptional responses under TMV infection ad a salicylic treatment time course (2 and 6 hours). (B) Overlap between the trancriptome responses to TMV and Mpjaz1ko mutants with constitutively active jasmonate signalling (left), and treatment with the active dn-OPDA jasmonate (right). (C) Overlap between the trancriptome responses commonly found in TMV infection and the different hormone-related transcriptomes. (D) Biological processes to which genes commonly upregulated in response to TMV infection, SA treatment (2,6 and 12h) and activation of jasmonate signalling belong (E) Biological processes to which genes commonly downregulated in response to TMV infection, SA treatment (12h) and activation of jasmonate signalling belong. (F) Biological processes to which commonly upregulated in response to TMV infection, SA treatment (6 h) and activation of jasmonate signalling in Mpjaz1KO plants belong. (G) Biological processes to which commonly upregulated in response to TMV infection, SA (6 h) and dn-OPDA treatments belong. Results in panels 4D-G come from a hypergeometric method with BY (BenjaminiYekutieli) p adjustment with minimal size of genes annotated for testing of 5 and p-value cutoff of 0.05.

To ascertain a possible contribution of the jasmonate signalling pathway, we compared the TMV transcriptional response with that of plants treated with dn-OPDA (Monte et al., 2018) or having a constitutively active jasmonate pathway as a consequence of the mutation of the JAZ signalling repressor (Mpjaz-1 KO plants; (Monte et al., 2019). A total of 143 and 81 genes were commonly up-regulated upon TMV infection and *MpJAZ* deficiency or dn-OPDA treatment, while 238 were downregulated in TMV infected and Mpjaz-1 KO plants (Figure 4B), suggesting that TMV infection also activates the jasmonate signalling pathway in Marchantia. Finally, we found that 64 genes were commonly up-regulated by TMV infection, SA 6 h treatment and absence of a functional MpJAZ protein, while 54 genes were up-regulated by TMV infection, SA 6h and dn-OPDA treatments (Figure 4C).

Next, we performed a GO analysis in the different sets of genes that were found to be commonly regulated by TMV infections and each of the different hormone-related transcriptomes (i.e. Up regulated genes by TMV and SA 6 h treatment from Figure 4A). That approach showed that the JA pathway might contribute to the activation of wounding, reactive oxygen species and virus responses, in addition to the catabolism of several compounds, including lipids, phenylpropanoids and salicylic acid, in TMV infected Marchantia plants. Interestingly, some of these processes were also commonly upregulated by TMV infection and SA treatment, including wounding, SA and lignin catabolism among others. The SA response pathway might on the other side also contribute to the observed transcriptional up-regulation of defense genes against bacteria and fungi, as well as response to chitin in TMV infected plants (Figure 4D, Supplementary Data 10). Amongst the downregulated processes by MpJAZ deficiency and TMV infection were translational elongation, mucilage and cell wall biosynthesis, response to fructose, photosynthesis, microtubule-based process, growth, and response to abscisic acid (ABA; Figure 4E, Supplementary Data 10)

Furthermore, genes found to be commonly upregulated by TMV, SA and dn-OPDA treatment were related to oxylipin and phenylalanine metabolism, wounding and drought recovery, among other Gene ontology groups (Figure 4F; Supplementary Data 11). Meanwhile, those genes found to be commonly upregulated by TMV infection, SA 6 h treatment and enhanced JA signalling in Mpjaz-1 plants were related to antiviral defense and lipid catabolism, which has been also recently found to contribute to SA activation and antiviral defense (Xu et al., 2022); Figure 4G, Supplementary Data 11).

Collectively, those results suggest that SA and JA signalling pathways are activated in Marchantia during TMV infection.

### TMV infection triggers aging in *Marchantia polymorpha*

Interactions with pathogens, including viruses, can result in the modification of life-history traits and developmental trajectories both in plants and animal (Pagan et al., 2008, Sievert, 1978). Senescence is the final stage of the life cycle of a cell, tissue or organism and it is gradually triggered throughout aging. This process can be accelerated by the interaction with different pathogens, including the crucifer infecting TMV-Cg strain capable of compatible interactions with the Arabidopsis UK-4 ecotype (Espinoza et al., 2007, Lim and Nam, 2005). Wound response, mitotic cell arrest, and reduced photosynthetic efficiency are hallmarks for plant cell senescence (Espinoza et al., 2007, Gan, 2003). Additionally, SA and JA, along with ET, contribute to senescence in *Arabidopsis thaliana* (Schommer et al., 2008, Yu et al., 2021). Since all those senescence hallmarks were present in our molecular characterization of the Marchantia-TMV interaction, we first checked the expression levels of several senescence marker genes in our transcriptomic data (Kulshrestha et al., 2022). We found that many senescence markers, such as MpMYB14 (Mp5g19050), MpORE1 (Mp2g07720), MpNYE1 (Mp1g17090) and MpWRKY7 (Mp3g17660), were induced upon TMV infection, hence suggesting that TMV infection could trigger aging and tissue maturation in Marchantia. Transcriptional trajectories orchestrating maturation and aging in Marchantia, both at organism and tissue level, have been recently reported to be very similar through transcriptional studies over developmental time, including individuals from different ages (spanning from 1 day after germination -D1- to 31 days old Marchantia plants -D31) , or by analysing sections along the thalli from top to base, being slice 1

(S1) at the top, considered the youngest part of the organ, and being slice 6 (S6) at the base and considered the oldest (Wang et al., 2023). Thus, while S1 samples were more similar to D1, those from S5 showed more commonalities to those from D31 plants. To assay whether TMV infection could modify the life-history of Marchantia plants we compared the transcriptional profile obtained from GFP positive areas from 4-week-old TMV-GFP infected Marchantia plants and that from equivalent zones from mock treated individuals with those from the maturation and developmental trajectories. Clustering of the transcriptional profile of mock and TMV infected plants with those datasets, based on Pearson’s correlation coefficient and regression model, showed that TMV infection promotes maturation and aging. Thus, while the transcriptome from mock treated plants clustered with juvenile state samples (slice 2 and Day 1), that from TMV infected plants clustered with slice 4 and day 31 of development (Figure 5A, 5B).

**Figure 5.**
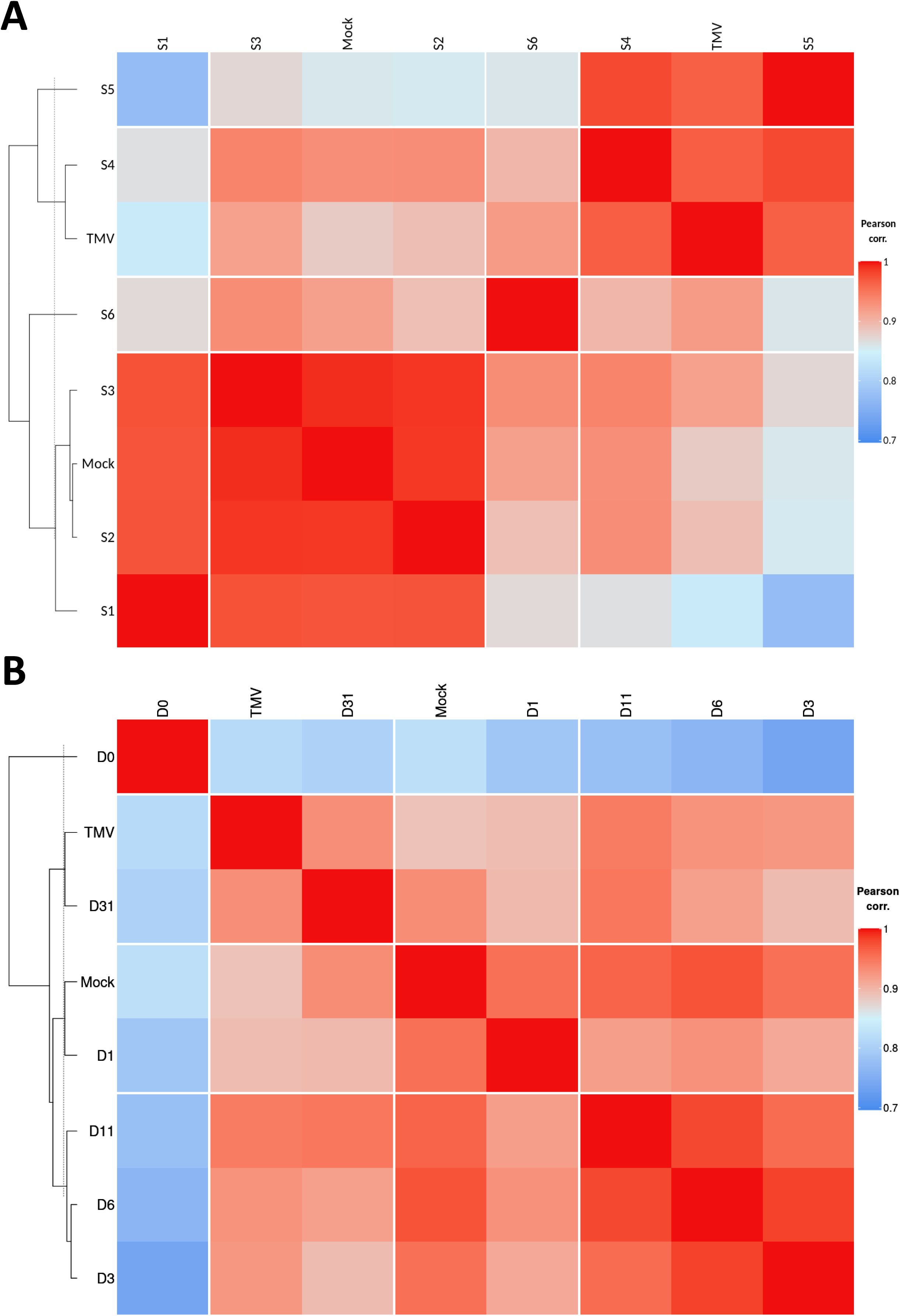
TMV infection triggers maturation and aging in Marchantia. (A) Pairwise comparison between transcriptomes from mock and TMV samples along with those from sliced thallus samples. Slice 1(S1) corresponds to the top of thalli (youngest part), while slice 6 (S6), corresponds to the base of thalli encompassing the oldest part. (B) Pairwise comparison between transcriptomes from mock and TMV samples along with those from Marchantia developmental series Day 1 (D1) samples correspond to 1 day after germination (youngest), while day 31 (31D) corresponds to fully mature Marchantia thalli 31 days after germination.

Those results indicated that TMV infection triggers premature tissue maturation and senescence in Marchantia, a characteristic trait from compatible plant - virus interactions.

### MpNAC7 is a conserved interactor of the TMV silencing suppressor p126 and enhances defense against TMV in tobacco plants

The TMV replicase p126 is essential for viral replication and movement, but it also works as a silencing suppressor in vascular plants (Ding et al., 2004). To gain insights into its possible role during TMV infection in Marchantia plants, we first established its subcellular localization. We found that p126 fused to GFP accumulated within the cytoplasm (Figure 6A) consistently with earlier studies in onion epidermal cells, where the full length p126 protein fused to GFP was found to be cytoplasmic (dos Reis Figueira et al., 2002).

**Figure 6.**
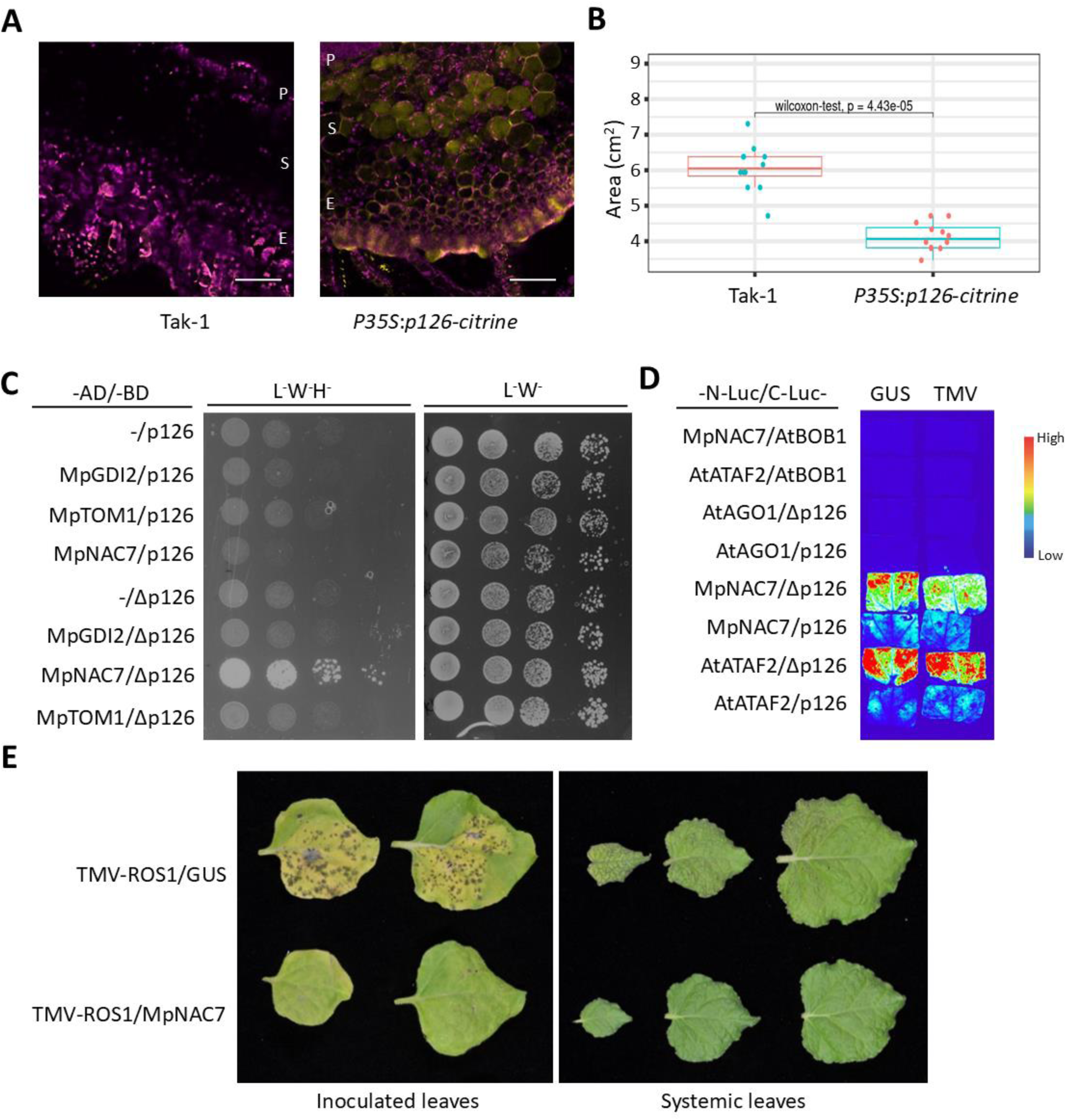
Conservation of the interaction between Marchantia and the TMV silencing suppressor p126. (A) p126-GFP shows accumulation in the cytoplasm of Marchantia cells in cross-sections of thalli from 4 weeks old plants (right) compared to thalli from untransformed plants of the same age plants (left). The pictures show the merge between green (GFP) and red (autofluorescence) channels. P stands for Photosynthetic region, S for storage region and E stands for lower epidermis. Merge was performed using Fiji Scale bars are 100 µMs. (B) Overexpression of p126 inhibits growth (area) when compared to Tak-1 control plants. Analysis was done in 4 weeks old plants in 3 biological replicates (n=12). Statistical significance was assessed using a two-sided Wilcoxon-test with Holm p adjustment. Length of the whiskers: 1.5 x IQR. Central line: median. Box width: IQR (i.e. Q3-Q1). (C) MpNAC7 interacts with the helicase domain of p126 in yeast. (D) MpNAC7 and ATAF2 interact both with the helicase and the full length p126 in planta. (E) Transient expression of MpNAC7 in Nicotiana plants abrogates local and systemic TMV infection. TMV presence is shown as purple.

Constitutive expression of viral-derived silencing suppressors often leads to plant developmental defects, like growth stunting (Kasschau et al., 2003). Accordingly, four-weeks old Marchantia plants constitutively expressing p126 were smaller in size when compared to control plants (Figure 6B, Supplementary Fig. 9).

The TMV replicase p126 is known to associate with several host factors in Arabidopsis, including TOM1 (Yamanaka et al., 2000), GDI2 (Kramer et al., 2011) and the transcription factor ATAF2 (Wang et al., 2009). To assess whether those interactions were conserved between p126 and the corresponding orthologs in Marchantia, we performed Y2H assays. Orthologs were identified by blasting the Arabidopsis protein sequences at MarpolBase (https//Marchantia.info) and PlantRegMap (http://plantregmap.gao-lab.org/). The p126 full length and helicase domain (Δp126, nt 2442 to 3636) were fused to the GAL4 binding domain and used as baits, while the putative interactors fused to the GAL4-activation domain, were used as preys. Notably, the only conserved interaction was that of the ortholog of the NAC transcription factor ATAF2, MpNAC7 (Mp6g02620), with Δp126 (Figure 6C). As for Δp126-ATAF2 binding (Wang et al., 2009) Δp126-MpNAC7 interaction could be confirmed *in planta* using bimolecular luminescence complementation (Figure 6D). Additionally, both the helicase domain alone and the full-length p126 were found to interact *in planta* with either NAC transcription factors. Such discrepancy between the results obtained by the two different protein-protein assays can be potentially attributed to the presence of additional protein partners in Nicotiana or variable conditions , such as intracellular pH or growth temperature, that modify protein folding allowing the interaction with the full length p126.

In Arabidopsis, ATAF2 expression is triggered by TMV infection (Wang et al., 2009). Likewise, we found that MpNAC7 expression is also increased on TMV infection (Log2(fold-change)= 0.991), wounding (Log2(fold-change)= 2.67), dn-OPDA and after 6 hours of SA treatment (Log2(fold-change)= 1,23) in Marchantia. Likewise, ATAF2 is induced by wounding, SA and JA treatments (Delessert et al., 2005) and contributes to defense against TMV in Arabidopsis (Wang et al., 2009). Here, to ascertain the role of MpNAC7 in defense against TMV, and rule out a possible link with RNA silencing-mediated responses, we infected 1 month old Nicotiana plants deficient in RDR6 (rdr6i; (Schwach et al., 2005)) with MpNAC7, or GUS as control, in the presence of TMV or an unrelated ssRNA virus, such as turnip mosaic virus (TuMV). TMV and TuMV strains were labelled with the ROS1 gene which triggers anthocyanin production, and hence we followed presence of the viruses by the purple colour (Bedoya et al., 2012). We observed that co-expression of MpNAC7 abrogated local and systemic TMV infection with respect to plants co-expressing the GUS control. By contrast, MpNAC7 co-expression was of no effect on TuMV infected plants, that displayed a similar local and systemic presence of the virus as the GUS co-infiltrated controls (Figure 6E, Supplementary Fig. 10).

Collectively, those results show that molecular interaction between the TMV silencing suppressor p126 and the host NAC transcription factor is conserved in Marchantia, as well as the role of this transcription factor in defense.

## Discussion

In this work, we have characterized the interaction of the non-vascular liverwort *Marchantia polymorpha* with viruses. First, we have defined known and unknown viruses that associate with Marchantia. This showed that Marchantia is primarily associated with RNA viruses in nature, validating previous predictions based on the presence of viral gene hallmarks in the transcriptomes from the 1000 Plants Genome Project (Mushegian et al., 2016). We also showed that at least one of these newly discovered viruses (Tübingen picorna-like virus 1) can replicate in laboratory grown Marchantia Tak-1 plants. In-depth phylogenetic analyses placed this virus within a larger, poorly characterised cluster composed of viruses discovered with metagenomics approaches, mostly associated with environmental samples, within the *Picornavirales* order. Viruses from this order infect mostly animals, but also protists and plants (Koonin et al., 2020) Tübingen picorna-like virus 1 codes for the hallmark protein domains characteristic of picornaviruses, but seems to lack movement protein (MP) that is present in most of the plant-infecting viruses, with the exception of some vertically transmitted viruses, e.g., from families *Paritiviridae* and *Amalgavirida (Dolja et al., 2020)*.Although kinetics of viral replication is not linear, in our infection experiment we observed only modest virus titre increase over time, which might be connected to limited ability of the virus to move from cell-to-cell due to the lack of MP. Few viruses have been identified by metagenomic approaches in non-vascular plants, such as *M. polymorpha*, some with (Debat et al., 2023, Mifsud et al., 2022) and other without MP (Mushegian et al., 2016, Vendrell-Mir et al., 2021). On the other hand, it is predicted that plasmodesmata have evolved along with the acquisition of vasculature to assist in the transport of nutrients throughout the plant by most likely changing the way they control their aperture (Benitez et al., 2018, Hernandez-Hernandez et al., 2020). Therefore, it is plausible that the way in which viruses interact with host plasmodesmata to move from cell-to-cell has also differed over time. Additionally, it has also been recently proposed that viruses lacking a MP can systemically infect vascular plants exploiting host proteins that are also present in Marchantia, such as those belonging to the Phloem Protein 2 (PP2) gene family (Ying et al., 2024). Since we only started to uncover the *M. polymorpha* virome, and, little is known about the structure and function of *M. polymorpha* plasmodesmata (Zeinsmaster D.D and Carothers Z.B, 1974), involved also in interaction with MP in vascular plants, further research building on the here developed *M. polymorpha* inoculation procedures, will contribute to shed a light on the repertoire of viruses infecting non-vascular plants. A recent study proposed a horizontal transfer of MP-encoding viruses infecting vascular plants to non-vascular plants (Butkovic et al., 2023), whereas discovery of Tübingen picorna-like virus 1 suggests another possible route of virus emergence in non-vascular plants. This may be linked with horizontal virus transfer from invertebrates or protists and could, very speculatively, represent a missing link in a recently suggested evolution of single jelly roll type CP to MP (Butkovic et al., 2023).

Second, we have found that TMV engages into a compatible interaction with Marchantia plants. That interaction encompasses an extensive transcriptional reprogramming reminiscent of that found in vascular plants such as Arabidopsis and Nicotiana, albeit involving also previously undescribed processes like the activation of a sustained wounding response here found in the Marchantia-TMV interaction. Third, we have found that the TMV p126 silencing suppressor displays similar subcellular localization in Marchantia cells, as in onion epidermal cells (dos Reis Figueira et al., 2002), and that it interacts with MpNAC7, a NAC transcription factor ortholog of the Arabidopsis ATAF2 factor involved in defense against TMV (Wang et al., 2009). Additionally, transient expression of MpNAC7 specifically abrogates TMV infection in Nicotiana plants. All those commonalities show that core molecular events of plant-virus interactions found in vascular plants might be also present in non-vascular plants.

TMV infected Marchantia plants undergo a transcriptional reprogramming consistent with a pathogenic process and characterized by the activation of defense related PR proteins and a conserved PTI response. Perception of dsRNAs has been found to trigger PTI in response to TMV infection in Arabidopsis, which in turn contributes to antiviral defense (Korner et al., 2013). Indeed, dsRNA-triggered PTI is dependent on the Arabidopsis PRR co-receptor SERK1 (Niehl et al., 2016), which has a corresponding ortholog in the Marchantia genome (Mp7g09160). This very same Marchantia gene has been suggested as the ortholog of the Arabidopsis BAK1 gene, which is central for PTI responses against *Pseudomonas syringae* (Gimenez-Ibanez, 2019). Mp7g09160 has been shown to interact with the *P. syringae* effector AvrPto, which abrogates PTI responses in Arabidopsis through interaction with AtBAK1 (Gimenez-Ibanez et al., 2019).. Thus, it will be interesting in future studies to analyse the role of Mp7g09160 in viral PTI responses. Additionally, TMV infection triggers the expression of a significant number of NLR genes that can potentially contribute to an ETI response. While in *Nicotiana benthamiana*, both DCL2 and DCL4 are processing TMV RNAs into 22 and 21 nucleotides long vsRNAs (Guo and Wong, 2017), the solely presence of DCL4 in Marchantia results in a predominant processing in 21 nucleotides long vsRNAs. We also observed changes on the expression of core genes of the RNA silencing machinery in response to TMV infection. In contrast to the repression of elements from the RNA-Directed DNA methylation pathway, such as MpDCL3, MpAGO9 and MpNRPE1b, the central elements within the RNA antiviral machinery MpDCL4 and MpRDR6 were activated. Such general switch on RNA silencing pathways is reminiscent of that found in *Nicotiana benthamiana* in response to TMV, although in *benthamiana* it is specifically restricted to the vasculature (Collum and Culver, 2017). These results suggest an evolutionary re-routing of defense mechanisms after acquisition of novel tissues or cell-types that become new focal points for plant-virus interactions.

Interestingly, our results also uncover a previously unknown role of the wound response as an antiviral defense mechanism. Plant cells undergoing wound responses experience inhibition of their photosynthetic ability becoming sink tissues for photoassimilates (Quilliam et al., 2006), attracting TMV movement (Brants, 1961). This wound response might be, at least partially, sustained by the activation of the salicylic and jasmonate pathways. Thus, it is tempting to speculate that sustained wound response in Marchantia plants, along with local and systemic decreased on photosynthetic capability, confine TMV infection limiting its movement throughout the plant. Additionally, premature senescence of infected tissues might further abrogate viral infection. Therefore, modification of host’s life-history traits, such as senescence, can constitute a defensive mechanism in Marchantia.

Salicylic and jasmonate pathways are central for defense against several pathogens. Although the interaction between both hormones has been traditionally considered antagonistic, recent studies suggest that such antagonism might have undergone evolutionary fine tuning, and hence depend on each plant-microbe interaction (de Vries et al., 2018). Antagonism of the salicylic and jasmonate pathways has been described in defense against bacterial and fungal pathogens in Marchantia (Gimenez-Ibanez, 2019; Matsui, 2020). Our comparative analysis of the transcriptional changes induced in response to TMV infection and those found in response to treatments with Salicylic and jasmonates (dn-OPDA), or in Mpjaz-1 KO plants, show that both SA and jasmonate pathways might contribute to the activation of genes involved in phenylpropanoid metabolism. Those findings are in line with Salicylic and jasmonates triggering this very same pathway in the moss *Physcomitrium patens,* which in turn is responsible for the production of defense-related compounds. (de Vries et al., 2018). Likewise, in line with their synergistic interaction being responsible for local and systemic resistance against TMV in *Nicotiana benthamiana* (Zhu et al., 2014), both salicylic and jasmonates were observed to activate genes involved in antiviral defense responses to TMV infection in Marchantia. Finally, a set of SA/dn-OPDA commonly upregulated genes might contribute to the above mentioned sustained wound response, and therefore, to contain virus spread.

Our work also highlights the conservation of central players, such as MpNAC7, in the specific defense against TMV. Future comparative research studies will help to ascertain the conservation of the genetic networks under MpNAC7 and ATAF2 regulation both in Marchantia and Arabidopsis.

Collectively, results from this work suggest the presence of evolutionary conserved and divergent plant strategies to cope with viral infections. Those might include gain and loss of defense strategies, such as those involving wound responses. Evolution of plant-virus interactions might also encompass the gradual acquisition of new elements within the underlying gene networks, as those physically interacting with pathogen-derived proteins. Finally, defense mechanisms might undergo anatomical re-location to newly pathogen-targeted cell-types and tissues over evolutionary times.

## Methods

### Plant growth conditions

Male *Marchantia polymorpha* plants Takaragaike-1 (Tak-1) were grown on plates with half strength Gamborǵs B5 medium (Duchefa) containing 1% agar and 1% sucrose under continuous light and 22°C.

*Nicotiana benthamiana* plants were grown on soil under long day conditions (16h light/ 8h dark) and 25°C.

Arabidopsis thaliana plants were grown on soil under long day conditions and 23 °C.

### Infection assays

Frozen tissue from Marchantia plants from the Tübingen population or lab grown Tak-1 plants was homogenized in liquid nitrogen cold mortars, resuspended in phosphate buffer pH7.2, filtered using 70 µM filters and kept at 4°C. Two-week old Tak-1 plants grown in sterile conditions in plates were vacuum incubated with each of both extracts and transferred to plates containing wet soil. Eight Tak-1 individuals exposed to each sap were collected either one or two weeks after incubation to assess viral replication.

Two week-old *Nicotiana benthamiana* plants were infiltrated with an agrobacterium solution (0.5 O.D) carrying the TMV-GFP clone (Peart et al., 2002). Infected Nicotiana plants were grown for 3 additional weeks along with non-infected plants and upper leaves displaying infection symptoms were collected, along with equivalent ones from non-infected plants and frozen in liquid nitrogen. Subsequently, leaf tissue was grinded in a liquid nitrogen cold mortar. Sap was isolated from grinded leaf tissue using phosphate buffer pH 7.2, filtered with 70 µM filters and maintained at 4°C. Two-weeks old Tak-1 plants were incubated either in TMV-GFP containing sap or sap from non-infected plants (mock) for 5 minutes in vacuum. After incubation, Tak-1 plants were transferred to plates containing wet soil and brought back to the growth chamber. GFP signal was monitored 1 and 2 weeks after incubation using a portable UV lamp or with a magnifying stereoscope couple to a UV light source.

### RNA isolation and expression analysis

Total RNA was extracted from Marchantia plants and Arabidopsis inflorescences using the protocol described in (Box et al., 2011). Marchantia isolated RNA was further cleaned using the RNeasy Kit (Qiagen). 500 ngs of total RNA were DNase I treated (Fisher Scientific) and converted into cDNA using the RevertAid first strand kit (Fisher Scientific) following the manufacturer instructions. RT-qPCR experiments were performed in a lightcycler (ROCHE) device. Relative expression changes were calculated using the 2^-ΔΔCt^ method using *MpACTIN7* (Mp6g11010) gene as housekeeping. Primers utilized can be found in supplementary table 3.

Samples for transcriptome assays were obtained from GFP positive areas collected from 5 4-week-old Marchantia plants per replicate (4 replicates total) after two weeks incubation with extracts from Nicotiana plants containing TMV-GFP. Equivalent areas were collected from mock treated 4-week-old Marchantia plants as control. RNA was

### RNA-seq analysis

For virome studies, 1 µg of total RNA from each of the 21 individuals included in the study was pooled and 4 µg from the mixture were used for preparing two independent libraries using total ribo-depleted RNA following the manufacturer instructions (Illumina TruSeq Stranded total RNA LT sample prep Kit, plant). The first library containing samples from 14 individuals was sequenced on an Illumina NovaSeq 6000 device in 2x150 bp mode, while the second one containing RNA from 7 individuals, was sequenced in an Illumina NovaseqX device. A total of 306,895,552 and 506,670,250 reads were respectively obtained. The reads were trimmed and further analysed using QIAGEN CLC Genomics Workbench 21 (https://digitalinsights.qiagen.com/). Dataset was analysed for the presence of known and new viruses using an established virus detection pipeline (Pecman et al., 2017), which includes: reads trimming, *de novo* assembly of trimmed reads, mapping of reads and contigs to NCBI viral RefSeq database (search for known viruses) and pFam domain search for contig sequences (search for known and unknown viruses). To detect known viruses, results of reads mapping to viral RefSeq database were inspected and filtered for viruses for which at least 1000 nts of the reference sequence were covered by mapped reads (Supplementary Data 1). In parallel, to detect known viruses and search for new ones, results of the pFam domain search of contig sequences were filtered for hits to viral proteins. Then, this candidate list of viral contigs was additionally queried for similarity against complete NCBI *nr* database (October 2020) using blastx (Gish and States, 1993) and the similarity search results were visualized and taxonomically classified using MEGAN 6 (Huson et al., 2016). Contigs corresponding to cellular organisms in this analysis were excluded from the list to obtain a final list of putative viral-like sequences (Supplementary Data 2). Trimmed sequencing reads were mapped to the list of putative viral-like sequences, to estimate the abundance of corresponding nucleic acids in the sequenced sample and the results were reported as average mapping depth (number of times a position is covered by sequencing reads, averaged over a complete contig length; Supplementary Data 2).

The presence of selected viruses (three new and two known) was tested by RT-qPCR in RNA extracts of collected samples as described above and datamining approach was used to search for association of those viruses with different sample types linked with publicly available short read datasets. This was performed by querying Serratus infrastructure palmprint database (Edgar et al., 2022) by RdRp sequences of selected viruses using palmID tool (https://serratus.io/palmid).

For a newly discovered picorna-like virus, named Tübingen picorna-like virus 1, additional analyses were performed, aiming to ascertain its association with *M. polymorpha*. Putative protein domains were annotated on the viral genome by InterProScan search (https://www.ebi.ac.uk/interpro/search/sequence/, May 2024) of the two discovered open reading frames against all included databases. Amino acid sequence of ORF 1 of the virus (containing RNA-dependent polymerase (RdRp) domain) was queried for similarity against complete NCBI *nr* database (January 2022) using blastp. Fifteen best hits according to bit score values (which have at least 33% identity with the conserved part of the RdRp domain of the queried virus) were included into phylogenetic analysis. Alignment of the sequences of the conserved part of RdRp was performed using MAFFT 7.48 (Katoh and Standley, 2013), followed by alignment trimming using TrimAI 1.3 in Automated 1 mode (Capella-Gutiérrez et al., 2009). Then, best protein substitution model was selected using Model Finder (Kalyaanamoorthy et al., 2017) and maximum likelihood phylogenetic tree inference was performed based on the trimmed alignment using IQ-Tree 1.6.12 (Nguyen et al., 2015) using 1000 ultrafast bootstrap replicates (Hoang et al., 2018). The resulting phylogenetic tree was visualized using iTOL 6.5 (Letunic and Bork, 2021) and viral taxa on the tree were coloured according to the associated sample source, as designated in NCBI GenBank. To obtain even more in-depth information about the phylogenetic position of the newly discovered virus, its sequence was aligned also against members of the most related family-level cluster of RdRp sequences (f.0032) from RNA Viruses in Metatranscriptomes (RVMT) database (https://riboviria.org/) (Neri et al., 2022). Alignment and phylogenetic analyses were performed as described above, excluding TrimAI step.

For analysing the transcriptional response in Marchantia plants to TMV infection, 1 ug of total RNA was used for preparing each stranded library by mRNA enrichment following manufacturer indications (TrueSeq kit from Illumina). Libraries were sequenced on a Novaseq 6000 device. A total of 20-21 million pairs were obtained per sample. Adapter removal and trimming of low quality bases was done using Trim Galore! v0.6.1 (https://www.bioinformatics.babraham.ac.uk/projects/trim_galore/), removing reads shorter than 20 bps and non-paired reads after the trimming step. Ribosomal RNA reads were removed using SortMeRNA v2.1b (Kopylova et al., 2012), removing also reads that become unpaired after this filtering step.

Cleaned reads together with the transcriptome of *Marchantia polymorpha* were used to quantify gene expression at transcript level using Salmon v1.1.0 (Patro et al., 2017).

For this, the file MpTak1v5.1_r1.mrna.fasta containing the Marchantia transcriptome (v5.1) was downloaded from https://marchantia.info/download/tak1v5.1/ (as of 19-Jan-2021). This file was indexed using *Salmon index* and then used as input for *Salmon quant*, which was ran with parameters -l A, –validateMappings, --recoverOrphans, --rangeFactorizationBins 4, --seqBias, and –gcBias. Mapping rate was found to be in the 89.5%-92.58% range.

The R progam tximport v1.14.2 (Soneson et al., 2015) was used to aggregate Salmon’s transcript expression estimates at gene level and the *DESeqDataSetFromTximport* from the R package DESeq2 v1.26.0 (Love et al., 2014) was used to create an R object with count data and experimental information for all samples. A low-expression filter was then applied to remove any gene with less than 10 counts across all 8 analyzed samples, resulting in a final set of 14,980 genes (from 19,217 annotated genes). Then, the *DESeq* function, which, among other things, computes size factors for sample normalization, was applied to the filtered object. DESeq2’s *rlog* function was used to transform count data to the log2 scale while minimizing for differences between samples for low-expressed genes. The transformed data was fed to DESeq2’s *plotPCA* function to visualize a PCA plot of the samples.

The analysis of differential expression was done using DESeq2’s *results* function with the filtered and normalized object as input and parameter *alpha = 0.05*.

All datasets used in this study for comparative analysis (wound response and salicylic acid time course) were analysed in the very same way as described above.

All Gene ontology analysis (GO) were performed using PLAZA (https://bioinformatics.psb.ugent.be/plaza/). Redundancy on representative categories was reduced using Revigo (http://revigo.irb.hr/).

For correlation analysis of transcriptome responses between developmental trajectories and TMV response, raw reads were processed as described above, except that rRNA was removed using ribodetector_cpu v0.2.7 (Deng et al., 2022) and Salmon version was v1.9.0. Salmon’s counts were converted into a DESeq2 object and a low-expression filter was applied as described above. Next, the existing batch effect between time course samples and sliced tissue ones was removed using the Combat seq function from the sva package (Leek et al., 2023), v3.48.0. Then, values for biological replicates were averaged and Pearson correlation was computed for the resulting count matrix and displayed using the Heatmap function from the ComplexHeatmap package (Gu, 2022). The batch correction, correlation and heatmap plotting steps were based on the R script used in (Wang et al., 2023) and kindly provided by the authors.

### DNA-seq data mining

Data mining was utilized to search for possible DNA viruses associated with *M. polymorpha.* DNA-seq data from a recent study of *M. polymorpha* from natural populations (Beaulieu et al., 2023) was retrieved from the SRA database (BioProject PRJNA931118). 74 biosample accessions were selected to encompass diverse geographical locations and various *M. polymorpha* subspecies. Datasets were analyzed using a customized pipeline for virus detection, designed with the Snakemake workflow management system (v.7.32.4) (Köster et al., 2021). Some of the chosen accessions contained sequencing data from multiple runs, which were merged before analysis. The downloaded reads were processed using a custom script: (available at: https://github.com/LanaVogrinec/snakemake-marchantia). Briefly, reads were trimmed with trimmomatic (v.0.39) (Bolger et al., 2014) and assembled into contigs with MEGAHIT (v.1.2.9) (Li et al., 2015). A similarity search was performed against the NCBI *nr* database (February 2024) using DIAMOND (v.0.9.14) blastx (Buchfink et al., 2015) and the hits were taxonomically classified with MEGAN6 (v.6.25.9) (Huson et al., 2016). Contig sequences classified as viral (group ’Viruses’ in MEGAN analysis) were extracted and are available, together with their classifications, in Supplementary Data 6.

### sRNA library construction and data processing

TMV-GFP positive areas were collected from 2 Marchantia plants infected for 2 weeks in two biological replicates. RNA was isolated following the protocol described in (Box et al., 2011). 10 ngs of total RNA were used to build two independent libraries following manufacturer indications (RealSeq-AC, Somagenics). Libraries were sequenced in an Illumina NovaSeq device. A total of 26-35 million single-end reads (101 bp) were obtained per sample. Adapter removal and trimming of low quality bases was done using Trim Galore! v0.6.1 (https://www.bioinformatics.babraham.ac.uk/projects/trim_galore/) with parameters *-q 15 -- clip_R1 1 --length 16*.

Cleaned reads were then mapped to the TMV-GFP sequence using hisat2 (v2.1.0, (Kim et al., 2019)) with default parameters. The resulting sam files were sorted and indexed using Samtools (v1.15.1, (Danecek et al., 2021)) and the length distribution of reads mapping to the TMV-GFP sequence was computed using samtools stats.

### Constructs and plant transformation

cDNA from Marchantia mock treated plants was used to amplify the coding regions of *MpTOM1* (Mp3g00660), *MpGDI2* (Mp6g05010) and *MpNAC7* (Mp6g02620) by PCR using Phusion polymerase (NEB). Arabidopsis *Bobber1* (At5g53400) and *ATAF2* (At5g08790) were amplified using cDNA synthesized from RNA from Arabidopsis inflorescences. p126 and Δp126 were amplified using DNA from the TMV-GFP containing plasmid. All obtained fragments were further incubated with dATP and Taq DNA polymerase (NEB) for 1 hour before cloned into the gateway compatible vector PCR8/GW/TOPO TA (Invitrogen) or directly cloned to pENTR D-TOPO (Invitrogen) in the case of ATAF2. An entry vector containing the Arabidopsis AGO1 was kindly provided by Pablo Manavella. Inserts were cloned into their final vectors through LR reaction using LR clonase II (Invitrogen). All plasmids were submitted to Sanger sequencing to corroborate their integrity before used in the corresponding experiments. A list of primers used, and constructs can be found in supplementary table 3 and supplementary table 4.

Plant transformation was performed as described in (Ishizaki et al., 2008) and selected in Gamborǵs B5 media supplemented with Hygromycin 100mg/ml and Cefotaxime 100mg/ml.

### Histology and imaging

Marchantia plants were embedded in 8% agarose and cross-sections of around 200 µM were obtained using a vibratome. Confocal imaging was performed using Leica SP5. Images were processed using Fiji (Schindelin et al., 2012).

### Protein-protein interaction assays

Interactions in the yeast two-hybrid system were assayed on selective medium (L^-^, W^-^, H^-^). The assay was repeated 3 times. For the luciferase complementation assay, one-month old rdr6i Nicotiana plants were used. *Agrobacterium tumefaciens* cultures with the different constructs were mixed in the same amounts at a final OD600=0.6. Leaves were imaged three days after inoculation using a LAS4000 device. The assay was repeated 3 times with equivalent results.

### Photosynthetic measurements

10 measurements per plant were performed in 5 infected and 5 mock treated individuals using an IMAGING-PAM Maxi device (Heinz Walz GmbH). Quantification was done using the ImagingWing integrated software.

### Reporting summary

Further information on research design is available in the reporting summary linked to this article.

## Data availability

Raw data from the virome has been deposited in SRA and GenBank under Bioproject PRJNA1022866 (https://www.ncbi.nlm.nih.gov/bioproject/?term=PRJNA1022866) and BioSamples (SAMN37631837 and SAMN40978195). Raw data from TMV infected vs. Mock treated plants RNA-seq and TMV sRNA-seq have been deposit in SRA (Bioproject PRJNA1010197, https://www.ncbi.nlm.nih.gov/bioproject/?term=PRJNA1010197). Reconstructed genomic sequences of three novel viruses uncovered by virome analysis were deposited in NCBI GenBank (OR640980 https://www.ncbi.nlm.nih.gov/nuccore/OR640980, OR640981 https://www.ncbi.nlm.nih.gov/nuccore/OR640981, OR640982 https://www.ncbi.nlm.nih.gov/nuccore/OR640982). Raw data for other analysis included in this work can be found in the accompanying Source data file.

## Code availability

Original code for DNA virome analysis can be accessed at https://github.com/LanaVogrinec/snakemake-marchantia, which has deposited at ZENODO under the ID 10.5281/zenodo.13359141 (https://zenodo.org/records/13359141).

## Supporting information

Summary of Supplementary Data contents

## Acknowledgments

We would like to thank Juan Jose López-Moya, Miguel A. Blázquez and Salomé Prat for critical reading of the manuscript. Sebastian Schornack, Phil Carella, Alexander Summers, the CUBG curation team (Samuel Brockington and Mar Millan), Klaus Schlaeppi, Katja Rembold, Alba Gonzalez, Detlef Weigel, Alexandra Kehl, Brigitte Fiebig, Luisa Lanfranco, Mara Novero, Isabel Monte, Adrián Vojtassák, Sabine Zachgo and Claudia Gieshoidt for their help in obtaining Marchantia individuals for the virome assays. Neža Pajek Arambašić for help with pipeline automation. We would like to thank Eduardo Bejarano for providing the plasmid containing TMV-GFP and Antonio Darós for providing TuMV-ROS1 and TMV-ROS1 containing plasmids. Pablo Manavella for the AtAGO1-pEntr plasmid. Phil Carella for an updated annotation of NLR genes in Marchantia. We would also thank to Jia-Wei Wang, Long Wang and Jie Xu for sharing the code used for correlation analysis between TMV transcriptional responses and the different samples from Marchantia developmental trajectories.

The work in the MoRE lab is funded by RYC-2015-19154 (funded by MCIN/AEI/ 10.13039/501100011033 and by “ESF Investing in your future”), RTI2018-097262-B-I00 (funded by MCIN/AEI/ 10.13039/501100011033 and by “ERDF A way of making Europe”) and through the “Severo Ochoa Programme for Centres of Excellence in R&D” 2016-2019 (SEV-2015-0533) and 2020-2023 (CEX2019-000902-S) funded by MCIN/AEI/ 10.13039/501100011033,CSIC-MCIN (Grant 202240I201) and the CERCA and SGR (2021-SGR-00875) programmes from the Generalitat de Catalunya to I-R-S. T J-G was recipient of a Postdoctoral Fellowship from CRAG through the “Severo Ochoa Programme for Centres of Excellence in R&D” 2016-2019 (SEV-2015-0533). L. V-M was supported by BES-2016-076986 (funded by MCIN/AEI/ 10.13039/501100011033 and by “ESF Investing in your future”). The work at NIB was funded by Slovenian Research and Innovation Agency core and project financing P4-0407, P4-0165 and J4-4553.

## Authors contributions

I.R-S. and T.J-G. conceptualized the study and designed experiments. I.R-S. supervised the study and acquired funding. T.J-G. and I.R-S. isolated RNAs for virome analysis. D.K. and L.V. performed virome analysis. E.R-M. and L.V-M. performed expression analysis of presence of viruses in natural Marchantia populations and Marchantia infection with natural viruses. T.J-G. and I.R-S. performed histological and expression analysis of viral infections. I.R-S. and V.M.G-M. performed RNA isolation, sRNA libraries and analysis for transcriptomic assays. E.R-M. carried out all phenotypic and p126 histological studies. I.R-S. carried out the protein-protein interaction assays and performed infection assays in Nicotiana. I.R-S. and D.K. wrote the manuscript with input from all authors.

## References

Bally, J., Nakasugi, K., Jia, F., Jung, H., Ho, S. Y., Wong, M., Paul, C. M., Naim, F., Wood, C. C., Crowhurst, R. N., Hellens, R. P., Dale, J. L. & Waterhouse, P. M. 2015. The extremophile Nicotiana benthamiana has traded viral defence for early vigour. Nat Plants, 1, 15165.

Beaulieu, C., Libourel, C., MBADINGA Zamar, D. L., Mahboubi, K. E., Hoey, D. J., Keller, J., Girou, C., Clemente, H. S., Diop, I., Amblard, E., Théron, A., Cauet, S., Rodde, N., Zachgo, S., Halpape, W., Meierhenrich, A., Laker, B., Brautigam, A., Greiff, G. R., Szovenyi, P., Cheng, S., Tanizawa, Y., LEEBENS-Mack, J. H., Schmutz, J., Webber, J., Grimwood, J., Jacquet, C., Dunand, C., Nelson, J. M., Roux, F., Philippe, H., Schornack, S., Bonhomme, M. & Delaux, P.-M. 2023. The <MARCHANTIA> pangenome reveals ancient mechanisms of plant adaptation to the environment. bioRxiv, 2023.10.27.564390.

Beck, M., Wyrsch, I., Strutt, J., Wimalasekera, R., Webb, A., Boller, T. & Robatzek, S. 2014. Expression patterns of flagellin sensing 2 map to bacterial entry sites in plant shoots and roots. J Exp Bot, 65, 6487–98.

Bedoya, L. C., Martinez, F., Orzaez, D. & Daros, J. A. 2012. Visual tracking of plant virus infection and movement using a reporter MYB transcription factor that activates anthocyanin biosynthesis. Plant Physiol, 158, 1130–8.

Belanger, S., Zhan, J. & Meyers, B. C. 2023. Phylogenetic analyses of seven protein families refine the evolution of small RNA pathways in green plants. Plant Physiol, 192, 1183–1203.

Benitez, M., HERNANDEZ-Hernandez, V., Newman, S. A. & Niklas, K. J. 2018. Dynamical Patterning Modules, Biogeneric Materials, and the Evolution of Multicellular Plants. Front Plant Sci, 9, 871.

Bowles, A. M. C., Paps, J. & Bechtold, U. 2022. Water-related innovations in land plants evolved by different patterns of gene cooption and novelty. New Phytol, 235, 732–742.

Bowman, J. L., Kohchi, T., Yamato, K. T., Jenkins, J., Shu, S., Ishizaki, K., Yamaoka, S., Nishihama, R., Nakamura, Y., Berger, F., Adam, C., Aki, S. S., Althoff, F., Araki, T., ARTEAGA-Vazquez, M. A., Balasubrmanian, S., Barry, K., Bauer, D., Boehm, C. R., Briginshaw, L., CABALLERO-Perez, J., Catarino, B., Chen, F., Chiyoda, S., Chovatia, M., Davies, K. M., Delmans, M., Demura, T., Dierschke, T., Dolan, L., DORANTES-Acosta, A. E., Eklund, D. M., Florent, S. N., FLORES-Sandoval, E., Fujiyama, A., Fukuzawa, H., Galik, B., Grimanelli, D., Grimwood, J., Grossniklaus, U., Hamada, T., Haseloff, J., Hetherington, A. J., Higo, A., Hirakawa, Y., Hundley, H. N., Ikeda, Y., Inoue, K., Inoue, S. I., Ishida, S., Jia, Q., Kakita, M., Kanazawa, T., Kawai, Y., Kawashima, T., Kennedy, M., Kinose, K., Kinoshita, T., Kohara, Y., Koide, E., Komatsu, K., Kopischke, S., Kubo, M., Kyozuka, J., Lagercrantz, U., Lin, S. S., Lindquist, E., Lipzen, A. M., Lu, C. W., DE Luna, E., Martienssen, R. A., Minamino, N., Mizutani, M., Mizutani, M., Mochizuki, N., Monte, I., Mosher, R., Nagasaki, H., Nakagami, H., Naramoto, S., Nishitani, K., Ohtani, M., Okamoto, T., Okumura, M., Phillips, J., Pollak, B., Reinders, A., Rovekamp, M., Sano, R., Sawa, S., Schmid, M. W., Shirakawa, M., Solano, R., Spunde, A., Suetsugu, N., Sugano, S., Sugiyama, A., Sun, R., Suzuki, Y., Takenaka, M., et al. 2017. Insights into Land Plant Evolution Garnered from the Marchantia polymorpha Genome. Cell, 171, 287–304 e15

Box, M. S., Coustham, V., Dean, C. & Mylne, J. S. 2011. Protocol: A simple phenol-based method for 96-well extraction of high quality RNA from Arabidopsis. Plant Methods, 7, 7.

Brants, D. H. 1961. The Influence of Meristematic Tissue and Injuries on the Transport of Tobacco Mosaic Virus in Nicotiana tabacum L. cultivar. Samsun. Acta Botanica Neerlandica, 10, 113–163.

Butkovic, A., Dolja, V. V., Koonin, E. V. & Krupovic, M. 2023. Plant virus movement proteins originated from jelly-roll capsid proteins. PLoS Biol, 21, e3002157.

Campbell, R. N. 1996. Fungal transmission of plant viruses. Annu Rev Phytopathol, 34, 87–108.

CAPELLA-Gutiérrez, S., SILLA-Martínez, J. M. & Gabaldón, T. 2009. trimAl: a tool for automated alignment trimming in large-scale phylogenetic analyses. Bioinformatics, 25, 1972–1973.

Carella, P., Gogleva, A., Hoey, D. J., Bridgen, A. J., Stolze, S. C., Nakagami, H. & Schornack, S. 2019. Conserved Biochemical Defenses Underpin Host Responses to Oomycete Infection in an Early-Divergent Land Plant Lineage. Curr Biol, 29, 2282–2294 e5.

Carella, P., Gogleva, A., Tomaselli, M., Alfs, C. & Schornack, S. 2018. Phytophthora palmivora establishes tissue-specific intracellular infection structures in the earliest divergent land plant lineage. Proc Natl Acad Sci U S A, 115, E3846–E3855.

Collum, T. D. & Culver, J. N. 2017. Tobacco mosaic virus infection disproportionately impacts phloem associated translatomes in Arabidopsis thaliana and Nicotiana benthamiana. Virology, 510, 76–89.

Chia, K. S., Kourelis, J., Teulet, A., Vickers, M., Sakai, T., Walker, J. F., Schornack, S., Kamoun, S. & Carella, P. 2024a. The N-terminal domains of NLR immune receptors exhibit structural and functional similarities across divergent plant lineages. Plant Cell, 36, 2491–2511.

Chia, K. S., Kourelis, J., Teulet, A., Vickers, M., Sakai, T., Walker, J. F., Schornack, S., Kamoun, S. & Carella, P. 2024b. The N-terminal domains of NLR immune receptors exhibit structural and functional similarities across divergent plant lineages. Plant Cell.

Danecek, P., Bonfield, J. K., Liddle, J., Marshall, J., Ohan, V., Pollard, M. O., Whitwham, A., Keane, T., Mccarthy, S. A., Davies, R. M. & Li, H. 2021. Twelve years of SAMtools and BCFtools. Gigascience, 10.

DE Vries, S., DE Vries, J., VON Dahlen, J. K., Gould, S. B., Archibald, J. M., Rose, L. E. & Slamovits, C. H. 2018. On plant defense signaling networks and early land plant evolution. Commun Integr Biol, 11, 1–14.

Debat, H., Garcia, M. L. & Bejerman, N. 2023. Expanding the Repertoire of the Plant-Infecting Ophioviruses through Metatranscriptomics Data. Viruses, 15.

Delessert, C., Kazan, K., Wilson, I. W., VAN DER Straeten, D., Manners, J., Dennis, E. S. & Dolferus, R. 2005. The transcription factor ATAF2 represses the expression of pathogenesis-related genes in Arabidopsis. Plant J, 43, 745–57.

Deng, Z. L., Munch, P. C., Mreches, R. & Mchardy, A. C. 2022. Rapid and accurate identification of ribosomal RNA sequences via deep learning. Nucleic Acids Res, 50, e60.

Ding, S. W., Han, Q., Wang, J. & Li, W. X. 2018. Antiviral RNA interference in mammals. Curr Opin Immunol, 54, 109–114.

Ding, S. W. & Voinnet, O. 2007. Antiviral immunity directed by small RNAs. Cell, 130, 413–26.

Ding, X. S., Liu, J., Cheng, N. H., Folimonov, A., Hou, Y. M., Bao, Y., Katagi, C., Carter, S. A. & Nelson, R. S. 2004. The Tobacco mosaic virus 126-kDa protein associated with virus replication and movement suppresses RNA silencing. Mol Plant Microbe Interact, 17, 583–92.

Dolja, V. V., Krupovic, M. & Koonin, E. V. 2020. Deep Roots and Splendid Boughs of the Global Plant Virome. Annu Rev Phytopathol, 58, 23–53.

DOS REIS Figueira, A., Golem, S., Goregaoker, S. P. & Culver, J. N. 2002. A nuclear localization signal and a membrane association domain contribute to the cellular localization of the Tobacco mosaic virus 126-kDa replicase protein. Virology, 301, 81–9.

Edgar, R. C., Taylor, B., Lin, V., Altman, T., Barbera, P., Meleshko, D., Lohr, D., Novakovsky, G., Buchfink, B., AL-Shayeb, B., Banfield, J. F., DE LA Pena, M., Korobeynikov, A., Chikhi, R. & Babaian, A. 2022. Petabase-scale sequence alignment catalyses viral discovery. Nature, 602, 142–147.

Erickson, F. L., Holzberg, S., CALDERON-Urrea, A., Handley, V., Axtell, M., Corr, C. & Baker, B. 1999. The helicase domain of the TMV replicase proteins induces the N-mediated defence response in tobacco. Plant J, 18, 67–75.

Espinoza, C., Medina, C., Somerville, S. & ARCE-Johnson, P. 2007. Senescence-associated genes induced during compatible viral interactions with grapevine and Arabidopsis. J Exp Bot, 58, 3197–212.

Gan, S. 2003. Mitotic and postmitotic senescence in plants. Sci Aging Knowledge Environ, 2003, RE7.

GARCIA-Ruiz, H., Takeda, A., Chapman, E. J., Sullivan, C. M., Fahlgren, N., Brempelis, K. J. & Carrington, J. C. 2010. Arabidopsis RNA-dependent RNA polymerases and dicer-like proteins in antiviral defense and small interfering RNA biogenesis during Turnip Mosaic Virus infection. Plant Cell, 22, 481–96.

GIMENEZ-Ibanez, S., Zamarreno, A. M., GARCIA-Mina, J. M. & Solano, R. 2019. An Evolutionarily Ancient Immune System Governs the Interactions between Pseudomonas syringae and an Early-Diverging Land Plant Lineage. Curr Biol, 29, 2270–2281 e4.

Gish, W. & States, D. J. 1993. Identification of protein coding regions by database similarity search. Nat Genet, 3, 266–72.

Gu, Z. 2022. Complex Heatmap Visualization. iMeta.

Guo, S. & Wong, S. M. 2017. Deep sequencing analysis reveals a TMV mutant with a poly(A) tract reduces host defense responses in Nicotiana benthamiana. Virus Res, 239, 126–135.

HERNANDEZ-Hernandez, V., Benitez, M. & Boudaoud, A. 2020. Interplay between turgor pressure and plasmodesmata during plant development. J Exp Bot, 71, 768–777.

Heyman, J., Canher, B., Bisht, A., Christiaens, F. & DE Veylder, L. 2018. Emerging role of the plant ERF transcription factors in coordinating wound defense responses and repair. J Cell Sci, 131.

Hoang, D. T., Chernomor, O., VON Haeseler, A., Minh, B. Q. & Vinh, L. S. 2018. UFBoot2: Improving the Ultrafast Bootstrap Approximation. Molecular Biology and Evolution, 35, 518–522.

Huson, D. H., Beier, S., Flade, I., Górska, A., EL-Hadidi, M., Mitra, S., Ruscheweyh, H. J. & Tappu, R. 2016. MEGAN Community Edition - Interactive Exploration and Analysis of Large-Scale Microbiome Sequencing Data. Plos Computational Biology, 12.

Ishizaki, K., Chiyoda, S., Yamato, K. T. & Kohchi, T. 2008. Agrobacterium-mediated transformation of the haploid liverwort Marchantia polymorpha L., an emerging model for plant biology. Plant Cell Physiol, 49, 1084–91.

Iwama, R. E. & Moran, Y. 2023. Origins and diversification of animal innate immune responses against viral infections. Nat Ecol Evol, 7, 182–193.

Jeon, H.-W., Iwakawa, H., Maramoto, S., Herrfurth, C., Gustche, N., Schlüter, T., Kyozuka, J., Miyauchi, S., Feussner, I., Zachgo, S. & Nakagami, H. 2022. Contrasting and conserved roles of NPR pathways in diverged land plant lineages. bioRxiv.

Jia, X., Wang, L., Zhao, H., Zhang, Y., Chen, Z., Xu, L. & Yi, K. 2023. The origin and evolution of salicylic acid signaling and biosynthesis in plants. Mol Plant, 16, 245–259.

Kalyaanamoorthy, S., Minh, B. Q., Wong, T. K. F., VON Haeseler, A. & Jermiin, L. S. 2017. ModelFinder: fast model selection for accurate phylogenetic estimates. Nature Methods, 14, 587-+.

Kappagantu, M., Collum, T. D., Dardick, C. & Culver, J. N. 2020. Viral Hacks of the Plant Vasculature: The Role of Phloem Alterations in Systemic Virus Infection. Annu Rev Virol, 7, 351–370.

Kasschau, K. D., Xie, Z., Allen, E., Llave, C., Chapman, E. J., Krizan, K. A. & Carrington, J. C. 2003. P1/HC-Pro, a viral suppressor of RNA silencing, interferes with Arabidopsis development and miRNA unction. Dev Cell, 4, 205–17.

Katoh, K. & Standley, D. M. 2013. MAFFT Multiple Sequence Alignment Software Version 7: Improvements in Performance and Usability. Molecular Biology and Evolution, 30, 772–780.

Kim, D., Paggi, J. M., Park, C., Bennett, C. & Salzberg, S. L. 2019. Graph-based genome alignment and genotyping with HISAT2 and HISAT-genotype. Nat Biotechnol, 37, 907–915.

Kneeshaw, S., Soriano, G., Monte, I., Hamberg, M., Zamarreno, A. M., Garcia-Mina, J. M., Franco-Zorrilla, J. M., Kato, N., Ueda, M., Rey-Stolle, M. F., Barbas, C., Michavila, S., GIMENEZ-Ibanez, S., Jimenez-Aleman, G. H. & Solano, R. 2022. Ligand diversity contributes to the full activation of the jasmonate pathway in Marchantia polymorpha. Proc Natl Acad Sci U S A, 119, e2202930119.

Koonin, E. V., Dolja, V. V., Krupovic, M., Varsani, A., Wolf, Y. I., Yutin, N., Zerbini, F. M. & Kuhn, J. H. 2020. Global Organization and Proposed Megataxonomy of the Virus World. Microbiol Mol Biol Rev, 84.

Kopylova, E., Noé, L. & Touzet, H. 2012. SortMeRNA: fast and accurate filtering of ribosomal RNAs in metatranscriptomic data. Bioinformatics, 28, 3211–3217.

Korner, C. J., Klauser, D., Niehl, A., DOMINGUEZ-Ferreras, A., Chinchilla, D., Boller, T., Heinlein, M. & Hann, D. R. 2013. The immunity regulator BAK1 contributes to resistance against diverse RNA viruses. Mol Plant Microbe Interact, 26, 1271–80.

Kramer, S. R., Goregaoker, S. P. & Culver, J. N. 2011. Association of the Tobacco mosaic virus 126kDa replication protein with a GDI protein affects host susceptibility. Virology, 414, 110–8.

Kulshrestha, S., Jibran, R., VAN Klink, J. W., Zhou, Y., Brummell, D. A., Albert, N. W., Schwinn, K. E., Chagne, D., Landi, M., Bowman, J. L. & Davies, K. M. 2022. Stress, senescence, and specialized metabolites in bryophytes. J Exp Bot, 73, 4396–4411.

Lee, W. S., Fu, S. F., Li, Z., Murphy, A. M., Dobson, E. A., Garland, L., Chaluvadi, S. R., Lewsey, M. G., Nelson, R. S. & Carr, J. P. 2016. Salicylic acid treatment and expression of an RNA-dependent RNA polymerase 1 transgene inhibit lethal symptoms and meristem invasion during tobacco mosaic virus infection in Nicotiana benthamiana. BMC Plant Biol, 16, 15.

Leek, J. T., Johnson, W. E., Parker, H. S., Fertig, E. J., Jaffe, A.E., Zhang, Y., Storey, J. D. & Torres, L. C. 2023. sva: Surrogate Variable Analysis.

Letunic, I. & Bork, P. 2021. Interactive Tree Of Life (iTOL) v5: an online tool for phylogenetic tree display and annotation. Nucleic Acids Research, 49, W293–W296.

Lewsey, M. G. & Carr, J. P. 2009. Effects of DICER-like proteins 2, 3 and 4 on cucumber mosaic virus and tobacco mosaic virus infections in salicylic acid-treated plants. J Gen Virol, 90, 3010–3014.

Liang, Y., Heyman, J., Xiang, Y., Vandendriessche, W., Canher, B., Goeminne, G. & DE Veylder, L. 2022. The wound-activated ERF15 transcription factor drives Marchantia polymorpha regeneration by activating an oxylipin biosynthesis feedback loop. Sci Adv, 8, eabo7737.

Lim, P. O. & Nam, H. G. 2005. The molecular and genetic control of leaf senescence and longevity in Arabidopsis. Curr Top Dev Biol, 67, 49–83.

Liu, C. & Nelson, R. S. 2013. The cell biology of Tobacco mosaic virus replication and movement. Front Plant Sci, 4, 12.

Liu, Y., Gao, Q., Wu, B., Ai, T. & Guo, X. 2009. Ngrdr1, an RNA-dependent RNA polymerase isolated from Nicotiana glutinosa, was involved in biotic and abiotic stresses. Plant Physiol Biochem, 47, 359–68.

LOPEZ-Gomollon, S. & Baulcombe, D. C. 2022. Roles of RNA silencing in viral and non-viral plant immunity and in the crosstalk between disease resistance systems. Nat Rev Mol Cell Biol, 23, 645–662.

Love, M. I., Huber, W. & Anders, S. 2014. Moderated estimation of fold change and dispersion for RNA-seq data with DESeq2. Genome Biol, 15, 550.

Lucas, W. J., Groover, A., Lichtenberger, R., Furuta, K., Yadav, S. R., Helariutta, Y., He, X. Q., Fukuda, H., Kang, J., Brady, S. M., Patrick, J. W., Sperry, J., Yoshida, A., LOPEZ-Millan, A. F., Grusak, M. A. & Kachroo, P. 2013. The plant vascular system: evolution, development and functions. J Integr Plant Biol, 55, 294–388.

Mandadi, K. K. & Scholthof, K. B. 2013. Plant immune responses against viruses: how does a virus cause disease? Plant Cell, 25, 1489–505.

Matsui, H., Iwakawa, H., Hyon, G. S., Yotsui, I., Katou, S., Monte, I., Nishihama, R., Franzen, R., Solano, R. & Nakagami, H. 2020. Isolation of Natural Fungal Pathogens from Marchantia polymorpha Reveals Antagonism between Salicylic Acid and Jasmonate during Liverwort-Fungus Interactions. Plant Cell Physiol, 61, 265–275.

Mifsud, J. C. O., Gallagher, R. V., Holmes, E. C. & Geoghegan, J. L. 2022. Transcriptome Mining Expands Knowledge of RNA Viruses across the Plant Kingdom. J Virol, 96, e0026022.

Monte, I., Franco-Zorrilla, J. M., Garcia-Casado, G., Zamarreno, A. M., Garcia-Mina, J. M., Nishihama, R., Kohchi, T. & Solano, R. 2019. A Single JAZ Repressor Controls the Jasmonate Pathway in Marchantia polymorpha. Mol Plant, 12, 185–198.

Monte, I., Ishida, S., Zamarreno, A. M., Hamberg, M., Franco-Zorrilla, J. M., Garcia-Casado, G., Gouhier-Darimont, C., Reymond, P., Takahashi, K., Garcia-Mina, J. M., Nishihama, R., Kohchi, T. & Solano, R. 2018. Ligand-receptor co-evolution shaped the jasmonate pathway in land plants. Nat Chem Biol, 14, 480–488.

Mushegian, A., Shipunov, A. & Elena, S. F. 2016. Changes in the composition of the RNA virome mark evolutionary transitions in green plants. BMC Biol, 14, 68.

Neri, U., Wolf, Y. I., Roux, S., Camargo, A. P., Lee, B., Kazlauskas, D., Chen, I. M., Ivanova, N., ZEIGLER Allen, L., PAEZ-Espino, D., Bryant, D. A., Bhaya, D., Consortium, R. N. A. V. D., Krupovic, M., Dolja, V. V., Kyrpides, N. C., Koonin, E. V. & Gophna, U. 2022. Expansion of the global RNA virome reveals diverse clades of bacteriophages. Cell, 185, 4023–4037 e18.

Nguyen, L. T., Schmidt, H. A., VON Haeseler, A. & Minh, B. Q. 2015. IQ-TREE: A Fast and Effective Stochastic Algorithm for Estimating Maximum-Likelihood Phylogenies. Molecular Biology and Evolution, 32, 268–274.

Niehl, A., Wyrsch, I., Boller, T. & Heinlein, M. 2016. Double-stranded RNAs induce a pattern-triggered immune signaling pathway in plants. New Phytol, 211, 1008–19.

Pagan, I., Alonso-Blanco, C. & Garcia-Arenal, F. 2008. Host responses in life-history traits and tolerance to virus infection in Arabidopsis thaliana. PLoS Pathog, 4, e1000124.

Patro, R., Duggal, G., Love, M. I., Irizarry, R. A. & Kingsford, C. 2017. Salmon provides fast and bias-aware quantification of transcript expression. Nature Methods, 14, 417-+.

Peart, J. R., Cook, G., Feys, B. J., Parker, J. E. & Baulcombe, D. C. 2002. An EDS1 orthologue is required for N-mediated resistance against tobacco mosaic virus. Plant J, 29, 569–79.

Pruitt, R. N., Locci, F., Wanke, F., Zhang, L., Saile, S. C., Joe, A., Karelina, D., Hua, C., Frohlich, K., Wan, W. L., Hu, M., Rao, S., Stolze, S. C., Harzen, A., Gust, A. A., Harter, K., Joosten, M., Thomma, B., Zhou, J. M., Dangl, J. L., Weigel, D., Nakagami, H., Oecking, C., Kasmi, F. E., Parker, J. E. & Nurnberger, T. 2021. The EDS1-PAD4-ADR1 node mediates Arabidopsis pattern-triggered immunity. Nature, 598, 495–499.

Quilliam, R. S., Swarbrick, P. J., Scholes, J. D. & Rolfe, S. A. 2006. Imaging photosynthesis in wounded leaves of Arabidopsis thaliana. J Exp Bot, 57, 55–69.

Rakhshandehroo, F., Rezaee, S. & Palukaitis, P. 2017. Silencing the tobacco gene for RNA-dependent RNA polymerase 1 and infection by potato virus Y cause remodeling of cellular organelles. Virology, 510, 127–136.

Redkar, A., GIMENEZ Ibanez, S., Sabale, M., Zechmann, B., Solano, R. & DI Pietro, A. 2022. Marchantia polymorpha model reveals conserved infection mechanisms in the vascular wilt fungal pathogen Fusarium oxysporum. New Phytol, 234, 227–241.

Schindelin, J., Arganda-Carreras, I., Frise, E., Kaynig, V., Longair, M., Pietzsch, T., Preibisch, S., Rueden, C., Saalfeld, S., Schmid, B., Tinevez, J. Y., White, D. J., Hartenstein, V., Eliceiri, K., Tomancak, P. & Cardona, A. 2012. Fiji: an open-source platform for biological-image analysis. Nat Methods, 9, 676–82.

Scholthof, K. B. 2004. Tobacco mosaic virus: a model system for plant biology. Annu Rev Phytopathol, 42, 13–34.

Schommer, C., Palatnik, J. F., Aggarwal, P., Chetelat, A., Cubas, P., Farmer, E. E., Nath, U. & Weigel, D. 2008. Control of jasmonate biosynthesis and senescence by miR319 targets. PLoS Biol, 6, e230.

Schwach, F., Vaistij, F. E., Jones, L. & Baulcombe, D. C. 2005. An RNA-dependent RNA polymerase prevents meristem invasion by potato virus X and is required for the activity but not the production of a systemic silencing signal. Plant Physiol, 138, 1842–52.

Sheng, Y., Yang, L., Li, C., Wang, Y. & Guo, H. 2019. Transcriptomic changes in Nicotiana tabacum leaves during mosaic virus infection. 3 Biotech, 9, 220.

Sievert, R.C.. 1978. Effect of Potato Virus Y and Tobacco Mosaic Virus on Field-Grown Burley Tobacco. Phytopathology, 68, 823–825.

Silvestri, A., Bansal, C. & Rubio-Somoza, I. 2024. After silencing suppression: miRNA targets strike back. Trends Plant Sci.

Soneson, C., Love, M. I. & Robinson, M. D. 2015. Differential analyses for RNA-seq: transcript-level estimates improve gene-level inferences. F1000Res, 4, 1521.

Vasseur, F., Baldrich, P., Jimenez-Gongora, T., Villar-Martin, L., Weigel, D. & Rubio-Somoza, I. 2022. miRNA472 deficiency enhances Arabidopsis thaliana defence without reducing seed production. bioRxiv, doi.org/10.1101/2022.12.13.520224

Vendrell-Mir, P., Perroud, P. F., Haas, F. B., Meyberg, R., Charlot, F., Rensing, S. A., Nogue, F. & Casacuberta, J. M. 2021. A vertically transmitted amalgavirus is present in certain accessions of the bryophyte Physcomitrium patens. Plant J, 108, 1786–1797.

Wang, L., Wan, M. C., Liao, R. Y., Xu, J., Xu, Z. G., Xue, H. C., Mai, Y. X. & Wang, J. W. 2023. The maturation and aging trajectory of Marchantia polymorpha at single-cell resolution. Dev Cell, 58, 1429–1444 e6.

Wang, X., Goregaoker, S. P. & Culver, J. N. 2009. Interaction of the Tobacco mosaic virus replicase protein with a NAC domain transcription factor is associated with the suppression of systemic host defenses. J Virol, 83, 9720–30.

Whitham, S., Dinesh-Kumar, S. P., Choi, D., Hehl, R., Corr, C. & Baker, B. 1994. The product of the tobacco mosaic virus resistance gene N: similarity to toll and the interleukin-1 receptor. Cell, 78, 1101–15.

Xu, T., Zhang, L., Zhen, J., Fan, Y., Zhang, C. & Wang, L. 2013. Expressional and regulatory characterization of Arabidopsis RNA-dependent RNA polymerase 1. Planta, 237, 1561–9.

Xu, X. J., Geng, C., Jiang, S. Y., Zhu, Q., Yan, Z. Y., Tian, Y. P. & Li, X. D. 2022. A maize triacylglycerol lipase inhibits sugarcane mosaic virus infection. Plant Physiol, 189, 754–771.

Yamanaka, T., Ohta, T., Takahashi, M., Meshi, T., Schmidt, R., Dean, C., Naito, S. & Ishikawa, M. 2000. Tom1, an Arabidopsis gene required for efficient multiplication of a tobamovirus, encodes a putative transmembrane protein. Proc Natl Acad Sci U S A, 97, 10107–12.

Ying, X. B., Dong, L., Zhu, H., Duan, C. G., Du, Q. S., Lv, D. Q., Fang, Y. Y., Garcia, J. A., Fang, R. X. & Guo, H. S. 2010. RNA-dependent RNA polymerase 1 from Nicotiana tabacum suppresses RNA silencing and enhances viral infection in Nicotiana benthamiana. Plant Cell, 22, 1358–72.

Yotsui, I., Matsui, H., Miyauchi, S., Iwakawa, H., Melkonian, K., Schluter, T., Michavila, S., Kanazawa, T., Nomura, Y., Stolze, S. C., Jeon, H. W., Yan, Y., Harzen, A., Sugano, S. S., Shirakawa, M., Nishihama, R., Ichihashi, Y., Ibanez, S. G., Shirasu, K., Ueda, T., Kohchi, T. & Nakagami, H. 2023. LysM-mediated signaling in Marchantia polymorpha highlights the conservation of pattern-triggered immunity in land plants. Curr Biol, 33, 3732–3746 e8.

Yu, D., Fan, B., Macfarlane, S. A. & Chen, Z. 2003. Analysis of the involvement of an inducible Arabidopsis RNA-dependent RNA polymerase in antiviral defense. Mol Plant Microbe Interact, 16, 206–16.

Yu, X., Xu, Y. & Yan, S. 2021. Salicylic acid and ethylene coordinately promote leaf senescence. J Integr Plant Biol, 63, 823–827.

Zamfir, A. D., Babalola, B. M., Fraile, A., Mcleish, M. J. & Garcia-Arenal, F. 2023. Tobamoviruses Show Broad Host Ranges and Little Genetic Diversity Among Four Habitat Types of a Heterogeneous Ecosystem. Phytopathology, 113, 1697–1707.

Zeinsmaster D.D & Carothers Z.B 1974. The Fine Structure of Oogenesis in Marchantia polymorpha. American Journal of Botany, 61, 499–512.

Zhu, F., Xi, D. H., Yuan, S., Xu, F., Zhang, D. W. & Lin, H. H. 2014. Salicylic acid and jasmonic acid are essential for systemic resistance against tobacco mosaic virus in Nicotiana benthamiana. Mol Plant Microbe Interact, 27, 567–77.

