## Supplementary figures and tables for "Conservation of molecular responses upon viral infection in the non-vascular plant *Marchantia polymorpha*"

**Supplementary Figures 1-10**

**Supplementary tables 1-4**

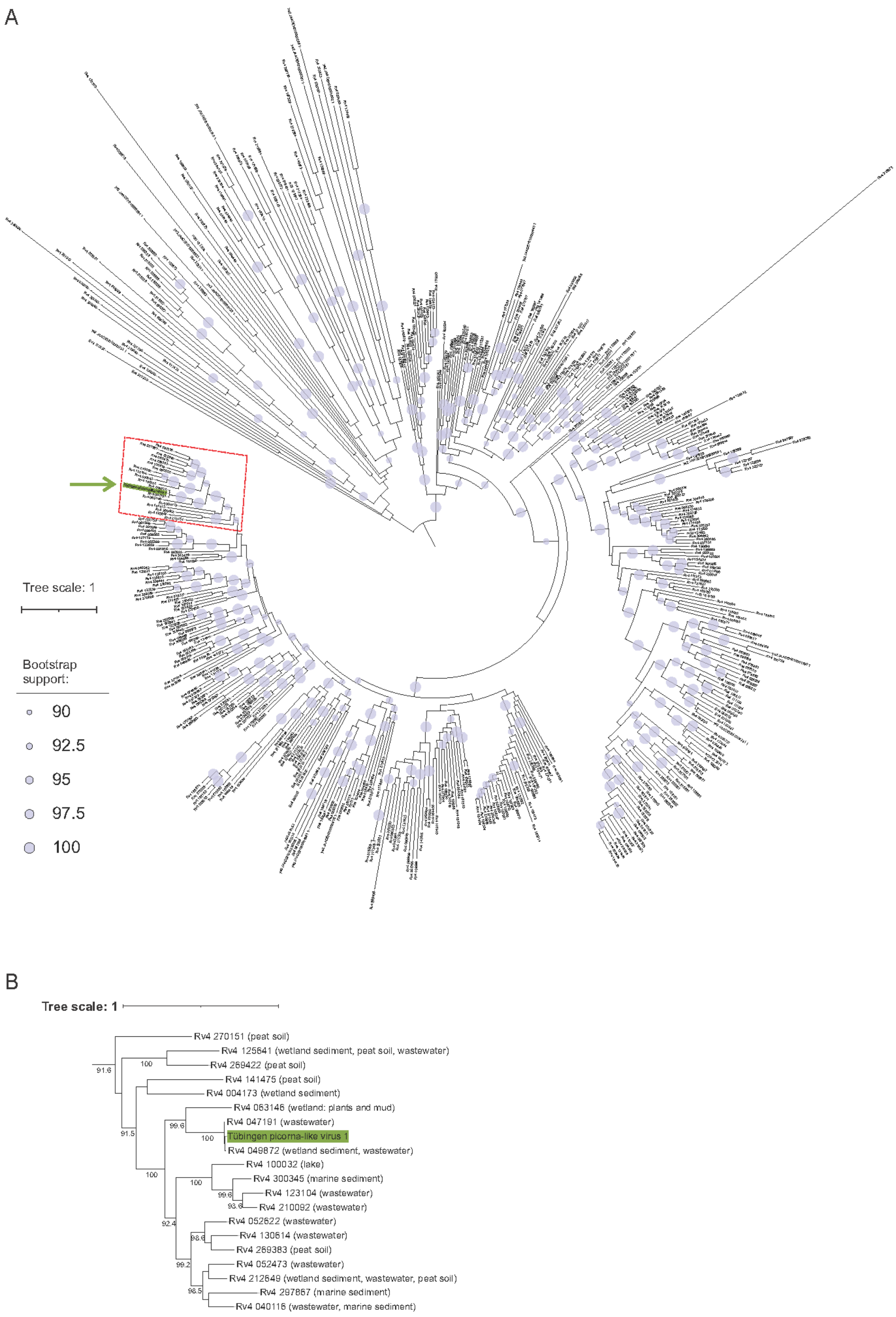
A
Tree scale: 1
Bootstrap support:
90
92.5
95
97.5
100
B
Tree scale: 1
Rv4 270151 (peat soil)
Rv4 125641 (wetland sediment, peat soil, wastewater)
Rv4 269422 (peat soil)
Rv4 141475 (peat soil)
Rv4 004173 (wetland sediment)
Rv4 063146 (wetland: plants and mud)
Rv4 047191 (wastewater)
Tübingen picorna-like virus 1
Rv4 049872 (wetland sediment, wastewater)
Rv4 100032 (lake)
Rv4 300345 (marine sediment)
Rv4 123104 (wastewater)
Rv4 210092 (wastewater)
Rv4 052622 (wastewater)
Rv4 130614 (wastewater)
Rv4 269383 (peat soil)
Rv4 052473 (wastewater)
Rv4 212649 (wetland sediment, wastewater, peat soil)
Rv4 297867 (marine sediment)
Rv4 040116 (wastewater, marine sediment)
91.6
100
91.5
99.6
100
100
100
99.6
93.6
92.4
98.6
99.2
98.5

**Supplementary figure 1**. **Phylogenetic position of a newly discovered Tübingen picorna-like virus 1 in regard to the most related RNA-dependent RNA polymerase (RdRp) sequences from RNA Viruses in Metatranscriptomes (RVMT) database (https://riboviria.org/).**

(A) Maximum likelihood phylogenetic tree obtained based on the alignment of the RdRd of Tübingen picorna-like virus 1 and 476 sequences corresponding to f.0032 cluster (within Picornavirales) from RVMT database. The tree was midpoint rooted. Tübingen picorna-like virus 1 is designated by green arrow. Ultrafast bootstrap support (>90%) is shown as circles according to the legend on the left. Branch length represents the average number of amino acid substitutions per site. Leaf labels represent RdRp sequences from RVMT database with additional info given in Supplementary table 5.

(B) A pruned phylogenetic tree from (A) showing in detail the clade containing RdRp sequences most similar to Tübingen picorna-like virus 1 (red box in (A)). Source sample environments linked with those RdRp sequences in RVMT database are shown in round brackets, with additional info given in

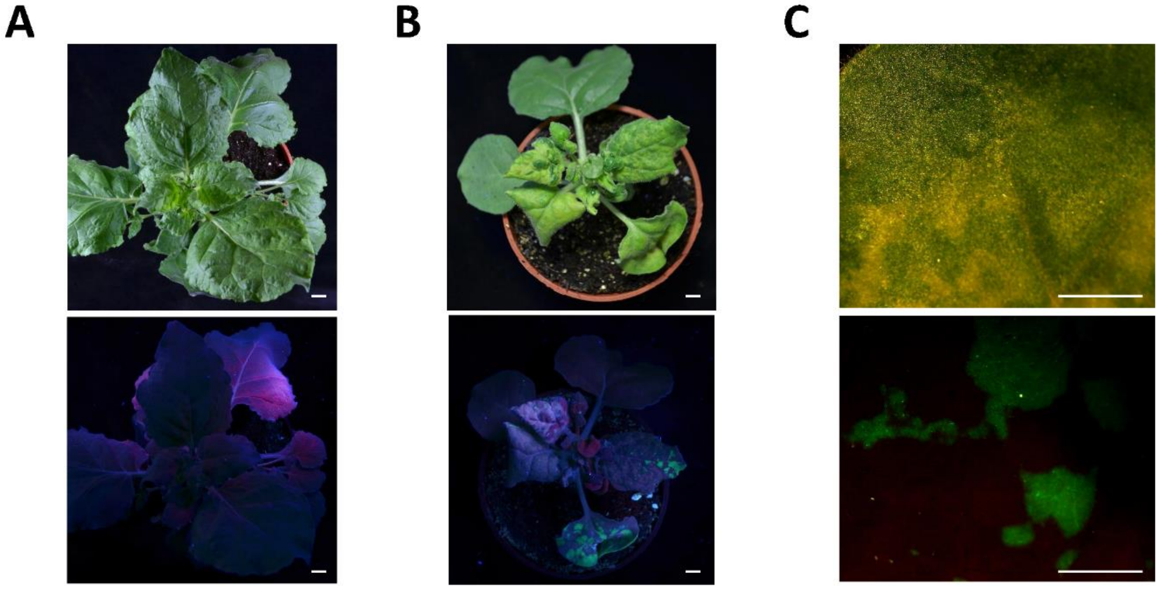


**Supplementary figure 2. Nicotiana benthamiana plants inoculated with sap from Marchantia infection experiments.**

(A) Nicotiana plants inoculated with sap from mock treated plants. Upper panel, top view of plant with white light. Down panel, same Nicotiana plant under UV light. Scale bars signify 1 cm.

(B) Nicotiana plants inoculated with sap from TMV-GFP treated plants. Upper panel, top view of plant with white light showing crinkled and yellowish leaves as result of TMV infection. Down panel, same Nicotiana plant under UV light showing GFP signal from TMV infection in leaves showing symptoms. Scale bars signify 1 cm.

[illegible]

**A**

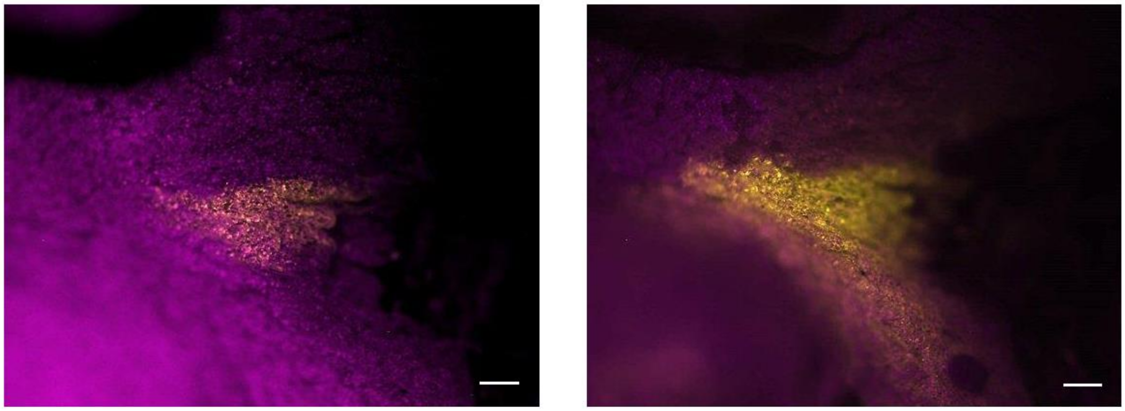

**Supplementary figure 3. TMV infections progress in Marhantia thalli over time.**

Thalli from Marchantia plants showing the progression of TMV infection inferred by GFP signal. Left, 3 weeks old plant after 1 week inoculation. Right, 4 weeks old plant after 2 weeks inoculation. Additional replicate to that shown in Figure 2A. Magenta shows autofluoresnce and yellow GFP signal.

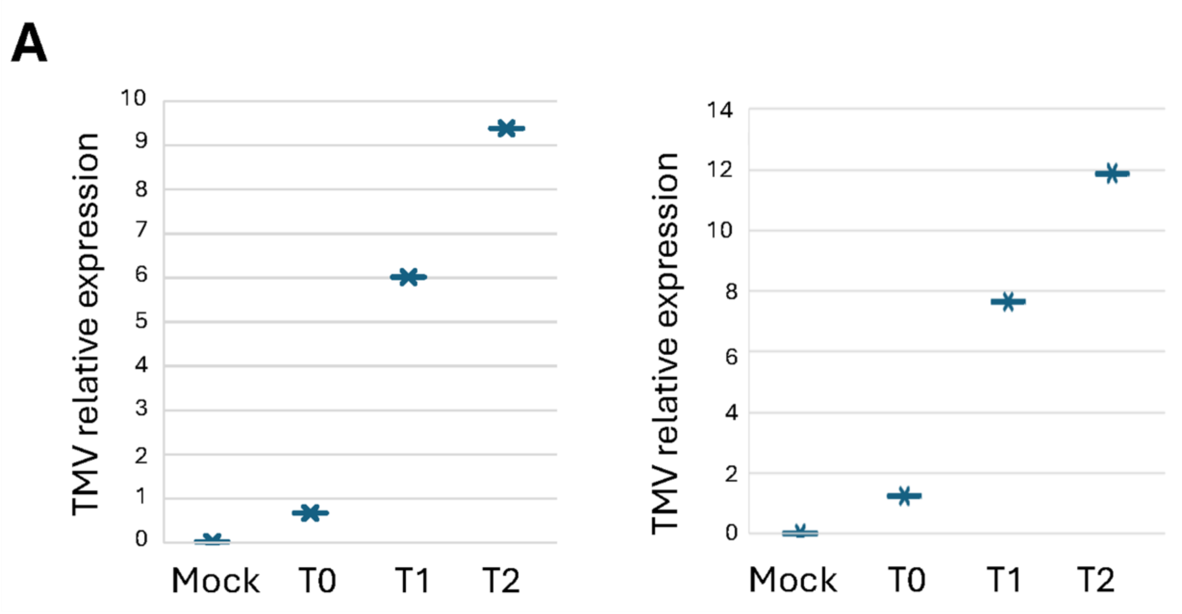


**Supplementary figure 4. TMV replicates in Marchantia plants.**

Two additional biological replicates to the one shown in Figure 2B. Whole individuals were collected in pools of 3 for each time point. Mock, 4 weeks old plants incubated with sap from untreated plants. T0, 2 weeks old plants collected right after incubation with sap from infected plants. T1, 3 weeks old plants collected after 1 week from incubation. T2, 4 weeks old plants collected after 3 weeks from incubation. Expression of TMV is relative to the expression of Marchantia actine.

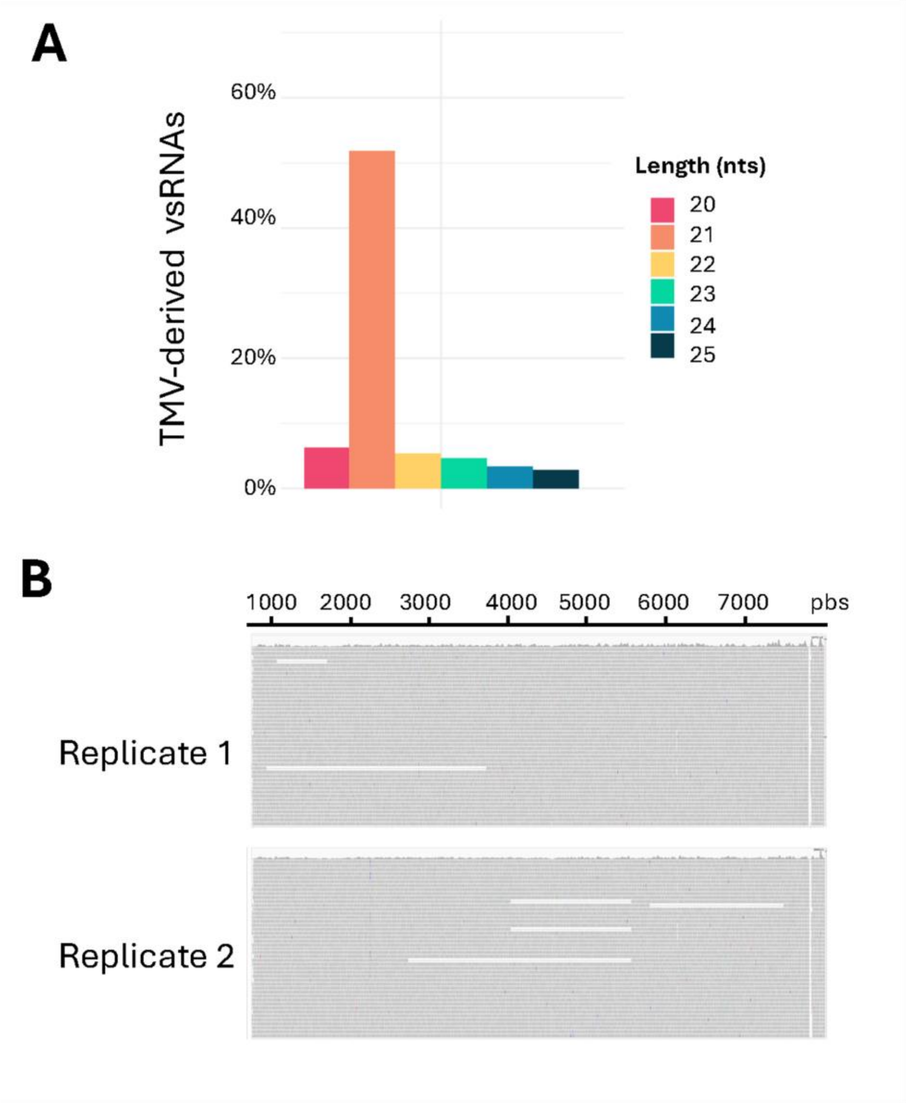


**Supplementary figure 5. Marchantia infected thalli accumulate TMV-derived vsRNAs.**

(A). Length distribution of TMV-derived vsRNAs profiled by sRNA sequencing showing that the predominant population belong to 21 nts long consistent with the absence of DCL2 and the presence of DCL4 in the Marchantia´s genome. Data shown belongs to replicate 2, being replicate 1 that shown in Figure 2B. Nts stans for nucleotides.

(B). Mapping of vsRNAs to the TMV-GFP sequence from both Replicate 1 (Figure 2B) and Replicate 2 (Sup. Figure 5A).

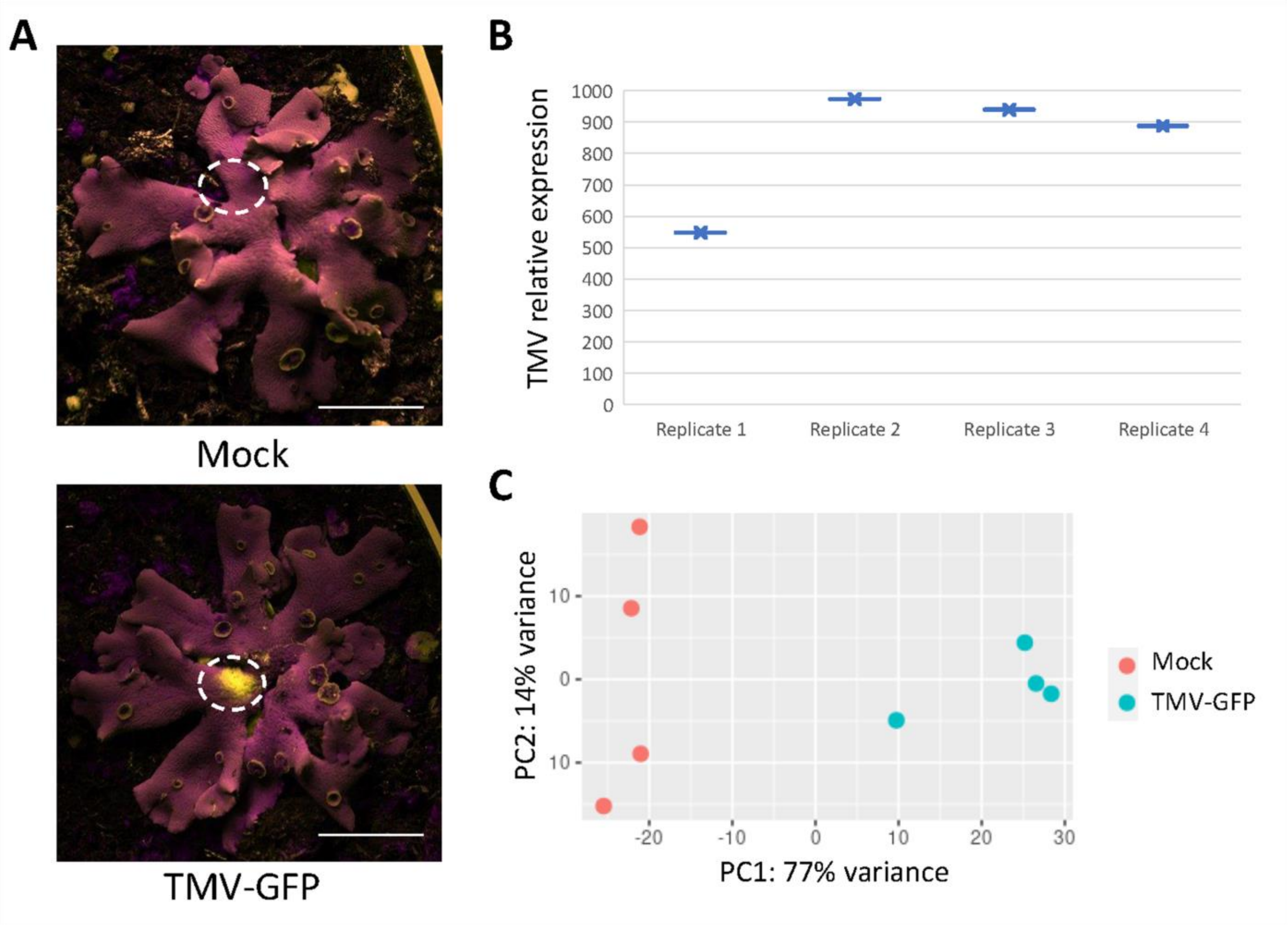


**Supplementary figure 6. Sampling and validation of samples for transcriptomic assays.**

(A) Example of areas collected from mock, upper panel, and TMV-GFP infected plants for transcriptomic assays. Scale bars signify 1 cm. Magenta shows autofluoresnce and yellow GFP signal.

(B) RT-qPCR validation of TMV-GFP levels in the 4 pools used for RNA-seq.

(C) Principal component analysis of the different replicates from RNA-seq experiment.

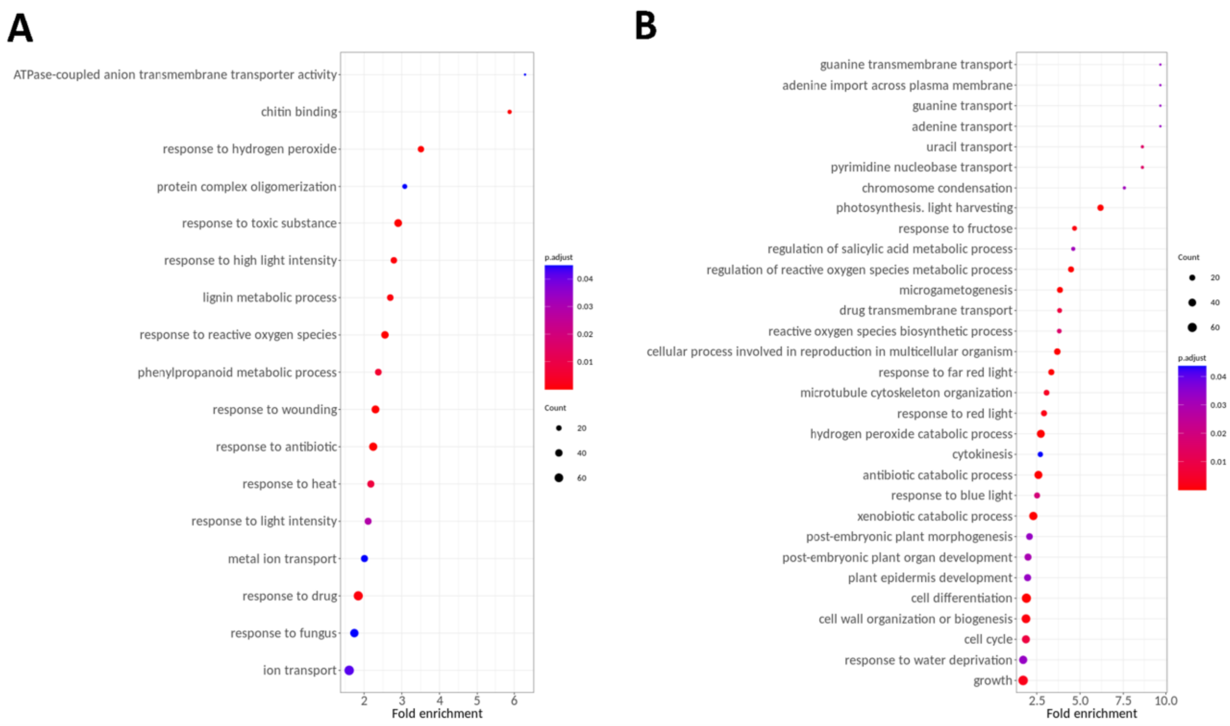


**Supplementary figure 7. Gene ontology analysis of biological processes to which DEGs in respond to TMV infection belong.**

(A) GO categories from upregulated genes in response to TMV infection.

(B) Selected GO categories from downregulated genes in response to TMV infection.

Results come from a hypergeometric method with BY (BenjaminiYekutieli) p adjustment with minimal size of genes annotated for testing of 5 and p-value cutoff of 0.05.

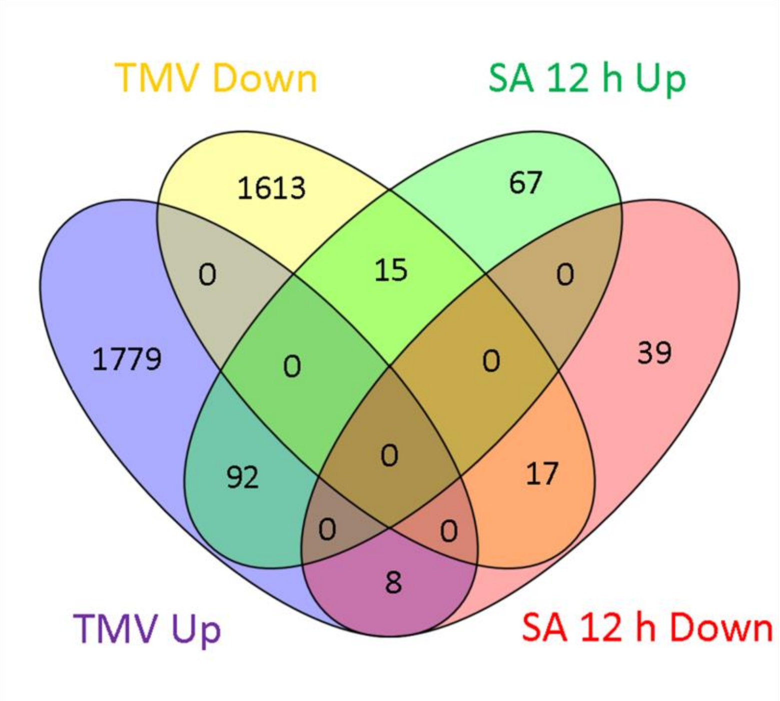


**Supplementary figure 8. Overlap between DEGs responding to 12 hours of incubation in salicylic acid and those responsive TMV infection.**

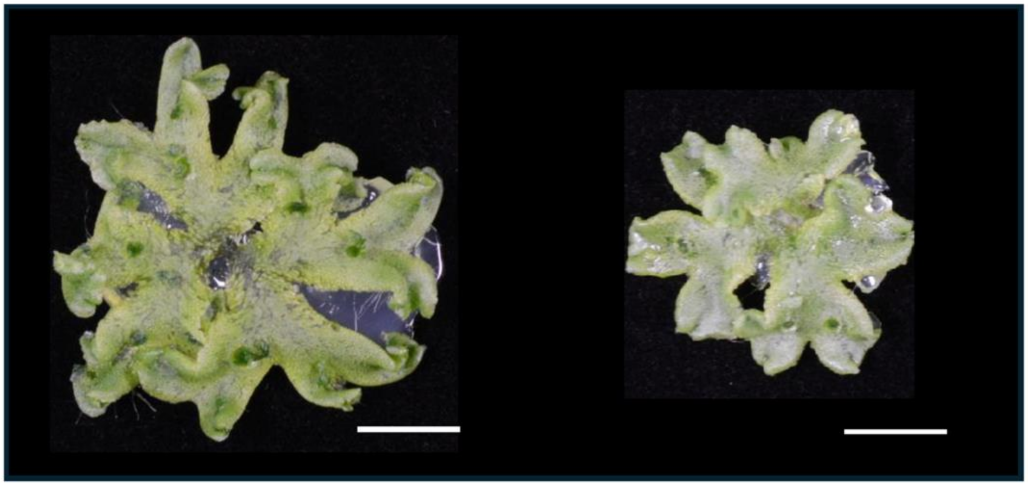


**Supplementary figure 9. p126 overexpression results in reduced growth.**

Representative 4 weeks old individuals from the phenotypic assay presented in Figure 6B. Scale bars represent 1 cm.

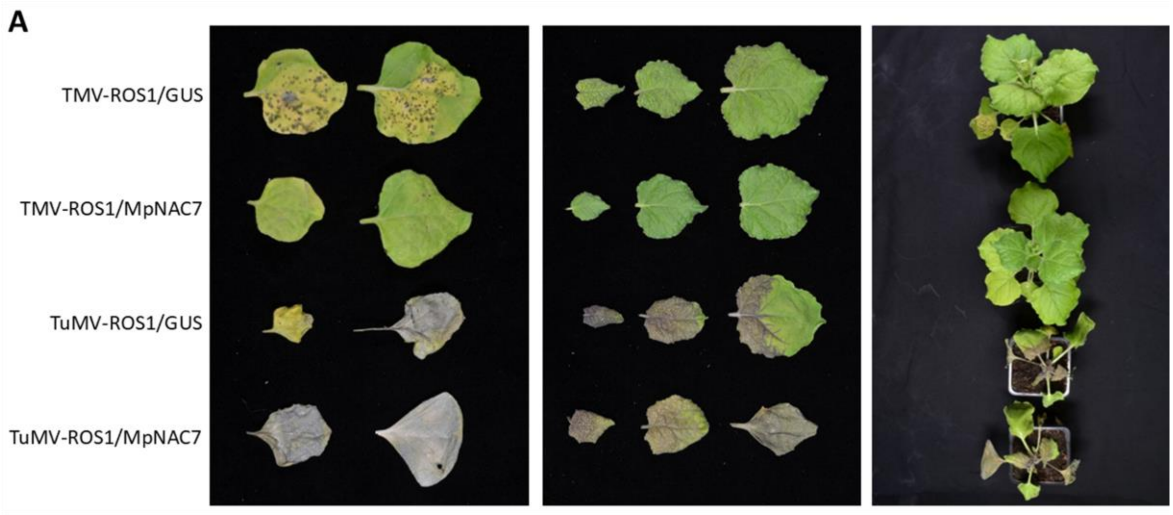


**Supplementary figure 10. MpNAC7 specifically abrogates TMV infection in Nicotiana plants.**

(A) Picture of inoculated leaves showing that MpNAC7 abrogates local TMV but not turnip mosaic virus (TuMV) infection (purple colour).

(B) Effect in systemic leaves.

(C) Upper view of infected plants.

| Name | ID | Relative expression level | padj |
|---|---|---|---|
| *MpPR1a* | Mp1g08990 | 2.96 | 0.0187 |
| *MpPR1d* | Mp2g06570 | 1.27 | 0.01 |
| *MpPR1o* | Mp5g16820 | 1.67 | 8.86E-08 |
| *MpPR1w* | Mp7g17200 | 1.74 | 1.82E-32 |
| *MpPR4* | Mp2g21910 | 2.99 | 0.0009 |
| *MpPR9* | Mp5g21510 | 4.04 | 1.18E-21 |
| *MpPR1e* | Mp2g11950 | -1.60 | 0.0244 |
| *MpPR1j* | Mp3g14130 | -2.97 | 0.0285 |
| *MpPR1m* | Mp5g12510 | -2.95 | 0.0088 |
| *MpPR1v* | Mp7g00440 | -2.78 | 5.68E-07 |
| *MpPR5a* | Mp1g15130 | -3.97 | 4.05E-22 |

**Supplementary table 1. List of Pathogenesis-related (PR) genes differentially expressed upon TMV infection.** DESeq2 results using the default parameters (Wald test and Benjamini-Hochberg p adjustment method)

| Name | ID | Relative expression level | p.ad |
|---|---|---|---|
| *MpNBS-LRR4* | Mp3g00170 | 1.7 | 2E-30 |
| *MpNBS-LRR6* | Mp3g09200 | 1.24 | 2E-09 |
| *MpNBS-LRR7* | Mp3g20180 | 1.84 | 1E-23 |
| *MpNBS-LRR10* | Mp4g08290 | 3.75 | 3E-83 |
| *MpNBS-LRR11* | Mp4g08790 | 6.6 | 7E-34 |
| *MpNBS-LRR12* | Mp4g21990 | 1.73 | 7E-26 |
| *MpNBS-LRR13* | Mp4g22030 | 1.75 | 2E-25 |
| *MpNBS-LRR17* | Mp5g21860 | 2.77 | 5E-18 |
| *MpNBS-LRR20* | Mp7g04670 | 1.42 | 3E-14 |

**Supplementary table 2. List of NLR genes differentially expressed upon TMV infection.** DESeq2 results using the default parameters (Wald test and Benjamini-Hochberg p adjustment method)

| Primer ID | Primer name | Sequence (5’-3’) | Purpose |
|---|---|---|---|
| M-693 | MpActine.s | AGGCATCTGGTATCCACGAG | RT-qPCR |
| M-694 | MpActine.as | ACATGGTCGTTCCTCCAGAC | RT-qPCR |
| M-1017 | TMV.s | GTAAGTTCCATGGGCCCTCCG | RT-qPCR |
| M-1018 | TMV.as | GGTTTGAGAGAGAAGATTACAAACGTG | RT-qPCR |
| M-913 | LNRV.s | GCAAGGCGACGATTGAGAAT | RT-qPCR |
| M-914 | LNRV.as | GAAGACCAGACCCACATCCA | RT-qPCR |
| M-915 | PLV.s | ACAAGGCCTTTCAAAGAGCG | RT-qPCR |
| M-916 | PLV.as | CAAAACCGCGTCTCCAAAGA | RT-qPCR |
| M-919 | TLV.s | TTTGCAAGAAGGTCCGTTGG | RT-qPCR |
| M-920 | TLV.as | TGCGCTGCCCATTACTTTAC | RT-qPCR |
| M-948 | Hubei.s | TGCCTCGTCTTCTTCACAGA | RT-qPCR |
| M-949 | Hubei.as | GCTCCAGTTATTTCCGGCTG | RT-qPCR |
| M-950 | SLV.s | AGCGGGAAAGTTTTGCTCAG | RT-qPCR |
| M-951 | SLV.as | CCCATGCGACAGAAATAGCC | RT-qPCR |
| M-833 | MpNAC7.s | ATGGTGATGGCGGCGAAGGG | Cloning |
| M-834 | MpNAC7.as | TCACCTTCCCCTTCCTGCGG | Cloning |
| M-858 | Mp.NAC7.ns.as | CCTTCCCCTTCCTGCGG | Cloning |
| M-835 | MpGDI2.s | ATGGATGAAGAGTATGATGTG | Cloning |
| M-836 | MpGDI2.as | TCACTCTTCTGCGGCACTAG | Cloning |
| M-841 | MpTOM1.s | ATGGATCCCGTTGTCGGATT | Cloning |
| M-842 | MpTOM1.as | TCATCGAATAGGATGGTATGCG | Cloning |
| M-715 | p126.s | ATGGCATACACACAGACAGC | Cloning |
| M-716 | p126.as | CTATTGTGTTCCTGCATCGACC | Cloning |
| M-717 | P126.ns.as | TTGTGTTCCTGCATCGACC | Cloning |
| M-1281 | ATAF2.s | CACCATGAAGTCGGAGCTAAATTTACC | Cloning |
| M-1282 | ATAF.ns.as | CCCCTGTGGAGCAAAACTCC | Cloning |

**Supplementary table 3. Table of primers used in this study.**

| Construct ID | Description | Vector | Used for |
|---|---|---|---|
| pN413 | P35S:p126-citrine | MpGWB106 | Transgenic line |
| pN363 | BD-p126/183 | pDest32 | Y2H assay |
| pN365 | AD-MpTOM1 | pDest22 | Y2H assay |
| pN369 | AD-MpNAC7 | pDest22 | Y2H assay |
| pN370 | AD-GDI1 | pDest22 | Y2H assay |
| pN407 | BD-Δp126/183 | pDest22 | Y2H assay |
| pN435 | C-Luc-p126/183 | pGW-C-Luc | BiLC assay |
| pN436 | C-Luc-Δp126/183 | pGW-C-Luc | BiLC assay |
| pN437 | MpNAC7-N-Luc | pGW-N-Luc | BiLC assay |
| pN440 | AtAGO1-N-Luc | pGW-N-Luc | BiLC assay |
| pN442 | C-Luc-AtBobber1 | pGW-C-Luc | BiLC assay |
| pN570 | ATAF2-N-Luc | pGW-N-Luc | BiLC assay |

**Supplementary table 4. Table of constructs used in this study.**
