## Supplementary material for "Conservation of molecular responses upon viral infection in the non-vascular plant *Marchantia polymorpha*": Summary of Supplementary Data contents

**Description of Additional Supplementary Files:**

**Supplementary Data 1.** Detection of viral sequences of known viruses.

**Supplementary Data 2.** Detection of known and putative novel virus-like sequences.

**Supplementary Data 3.** Demographic study for the presence of a set of known and newly identified virus in individuals from the different sampling sites.

**Supplementary Data 4.** Results of the palmID scan (https://serratus.io/palmid) for set of known and newly identified viruses.

**Supplementary Data 5.** Metadata for RdRp clusters showed on Supplementary figure 1.

**Supplementary Data 6.** Results of the mining of DNA sequencing data (BioProject PRJNA931118) for the presence of known and putative novel viral sequences.

**Supplementary Data 7.** List of DEG in response to TMV infection in Marchantia.

**Supplementary Data 8.** List of GO categories of the biological processes to which upregulated genes in response to TMV infection belong. Results come from a hypergeometric method with BY (BenjaminiYekutieli) p adjustment with minimal size of genes annotated for testing of 5 and p-value cutoff of 0.05.

**Supplementary Data 9.** List of GO categories of the biological processes to which downregulated genes in response to TMV infection belong. Results come from a hypergeometric method with BY (BenjaminiYekutieli) p adjustment with minimal size of genes annotated for testing of 5 and p-value cutoff of 0.05.

**Supplementary Data 10.** List of GO categories of the biological processes to which commonly responsive genes to TMV infection and salicylic acid time course treatment, Mpjaz-1ko mutants and dn-OPDA treatment belong. Results come from a hypergeometric method with BY (BenjaminiYekutieli) p adjustment with minimal size of genes annotated for testing of 5 and p-value cutoff of 0.05.

**Supplementary Data 11.** List of GO categories of the biological processes to which commonly responsive genes to TMV infection and different hormone-related transcriptomes belong. Results come from a hypergeometric method with BY (BenjaminiYekutieli) p adjustment with minimal size of genes annotated for testing of 5 and p-value cutoff of 0.05.
